# A *de novo* matrix for macroscopic living materials from bacteria

**DOI:** 10.1101/2021.11.12.468079

**Authors:** Sara Molinari, Robert F. Tesoriero, Dong Li, Swetha Sridhar, Rong Cai, Jayashree Soman, Kathleen R. Ryan, Paul D. Ashby, Caroline M. Ajo-Franklin

## Abstract

Engineered living materials (ELMs) are composites of living cells embedded in a biopolymer matrix that combine the desirable properties of natural biomaterials with non-natural, tailored properties. ELMs with a wide range of sophisticated biological functions have been created by engineering the embedded cells using synthetic biology. Engineering a *de novo* biomolecular matrix would offer control over material assembly, structure, and composition, thus enabling us to grow macroscopic ELMs with customizable mechanical properties. However, we have lacked the genetic tools and design rules to genetically encode a synthetic matrix that programs collective cell self-organization into macroscopic structures. Here we report growth of macroscopic ELMs from *Caulobacter crescentus* cells that display and secrete an engineered self-interacting protein. This protein formed an extracellular *de novo* matrix and assembled cells into hierarchically-ordered, centimeter-scale ELMs. We showed that the mechanical, catalytic, and morphological properties of these ELMs can be tuned through genetic modification of the matrix. Our work identifies novel genetic tools, design and assembly rules for growing macroscopic ELMs with both wide-ranging mechanical properties and customizable functions. We anticipate the modularity of this approach will permit the incorporation of different protein polymers in the *de novo* matrix, thus allowing to generate ELMs with a variety of desired structures and compositions of the bulk material. We envision specific matrix properties that can be combined synergistically with existing cellular functions to greatly expand the opportunities for ELMs in human health, energy, and the environment.

## Main Text

Naturally occurring living biomaterials, such as bones or wood, grow bottom-up from a small number of progenitor cells into macroscale structures^1^. Engineered living materials (ELMs)^2–4^ are inspired by naturally-occurring living materials, but use synthetic biology to introduce tailored, non-natural functions. By incorporating engineered cells into a biopolymer matrix, these materials can function as living sensors^5^, therapeutics^6^, biomanfacturing platforms^7^, electronics^8^, energy converters^9^, and structural materials^10^. While cells confer functionality to ELMs, the matrix assembles the material and controls the bulk material composition, structure, and function^11^.

Since the matrix plays such a key role over material properties, engineering macroscopic ELMs that grow from the bottom-up with a synthetic biomolecular matrix is a primary goal of the field. It is considered well beyond the current state-of-the art^11^ because secreting recombinant biopolymers at concentrations that gelate is challenging^12^ and because assembly of micrometer-sized cells into centimeter-scale materials requires self-organization across length scales spanning four orders of magnitude. Engineering principles to achieve this are unknown^11,12^, so most ELMs are microscopic^13–17^ and must be processed into macroscopic materials. The few macroscopic ELMs have been created by genetically modifying existing matrices^18^ or genetically manipulating mineralization of inorganic matrices^19^. However, these approaches have afforded little genetic control over the matrix composition and only ∼20-30% changes in material mechanics^18,19^, which is much more limited than the tunability of naturally-occurring and chemically synthesized materials.

### *De novo* engineering of a macroscopic bottom-up ELM

Leveraging previous genetic tools for biopolymer secretion and display in *Caulobacter crescentus*^20,21^, we sought to create bottom-up ELMs composed of cells interacting through a surface-bound *de novo* matrix. To minimize native cell-cell interactions, we started with a *C. crescentus* background that cannot form a biofilm. Next, we designed a <u>b</u>ottom-<u>u</u>p *<u>d</u>e novo* (BUD) protein by replacing the native copy of the surface layer (S-layer) RsaA^22^ (Fig. 1A) with a synthetic construct encoding four modules (Fig. 1B): (i) a surface-anchoring domain, (ii) a flexible biopolymer region for solution accessibility, (iii) a tag for functionalization, and (iv) a domain for secretion and self-interaction. We used the first 250 residues of RsaA as an anchor to the O-antigen lipopolysaccharide^23,24^. As the flexible domain, we chose an elastin-like polypeptide (ELP) based on human tropoelastin with 60 repeats of the Val-Pro-Gly-X-Gly motif^21,25^, ELP_60_. ELPs are flexible, self-associate, and form elastic structures. SpyTag^26^ was used as a functionalization tag, as it covalently binds to fusion proteins containing SpyCatcher. The C-terminal domain of the BUD protein, consisting of the last 336 residues of RsaA, was chosen to mediate protein secretion^21^ and to self-associate^27^. We refer to this BUD protein-expressing strain of *C. crescentus* as the BUD-ELM strain.

**Fig. 1.**
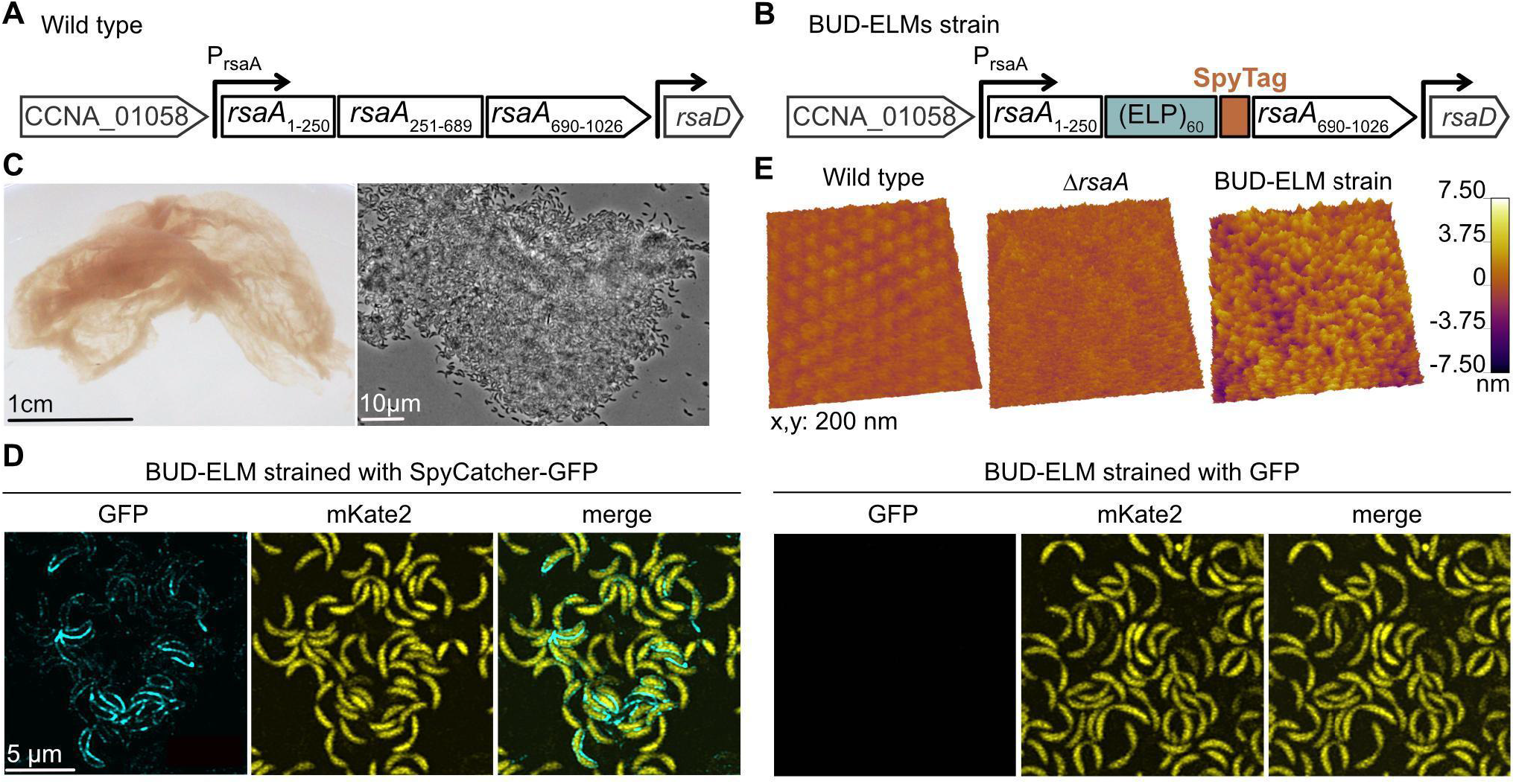
Engineered strains of *C. crescentus* self-assemble into BUD-ELMs. (**A**) Schematic of the native *rsaA* gene within its genomic context, showing its N-terminal cell anchoring domain (1-250) and C-terminal domain (251-1026) with the secretion subdomain (690-1026). (**B**) Schematic of the four-domain construct replacing the native *rsaA* gene in the BUD-ELM strain. (**C**) Photograph of free-floating material formed by the BUD-ELM strain (left). Brightfield image of a portion of a BUD-ELM (right), showing cell clusters and intact cells. (**D**) Confocal microscopy of single cells of BUD-ELM strain stained with SpyCatcher-GFP (left) or GFP (right), demonstrating that the BUD protein (Fig. 2A – left) is located on the cell surface. Scale bar: 5 µm, applies to all images. (**E**) AFM images of the cell surface of wild-type (left), *ΔrsaA* (middle), and BUD-ELM strain (right), showing the brush-like structure of the BUD-ELM strain’s surface.

Surprisingly, cultures of the BUD-ELM strain yielded centimeter-scale, filamentous material (Fig. 1C – left) after 24 h of growth. The material contained intact *C. crescentus* cells (Fig. 1C – right), indicating these macroscopic materials were indeed BUD-ELMs. In contrast, the wild-type culture did not generate any visible aggregates (Fig. S1). To understand the role of BUD protein in this material, we compared the extracellular surface of planktonic cells of the BUD-ELM strain before material formation with other *C. crescentus* strains. When stained with SpyCatcher-GFP, cells of the BUD-ELM strain (Fig. 1D – left) showed GFP fluorescence (cyan) along the outer contour of the cells (yellow), demonstrating the BUD protein is on the extracellular surface. The BUD-ELM was not stained by free GFP (Fig. 1D – right), nor was the Δ*SpyTag* strain stained by SpyCatcher-GFP (Fig. S2A and B), confirming that staining required SpyTag and SpyCatcher. Atomic force microscopy (AFM) of the BUD-ELM cells showed a brush-like structure (Fig 1E – right) that distinguished it from the wild-type hexameric S-layer (Fig. 1E – left) and the *ΔrsaA* strain (Fig. 1E – middle). The BUD protein formed long, unstructured projections, and this soft layer mediates cell-cell interactions (Fig. S3). These results provide a first demonstration of macroscopic, bottom-up ELMs with a *de novo* surface-bound matrix that mediates cell-cell interactions.

### BUD-ELMs are organized hierarchically through a synthetic proteinaceous matrix

To probe their structure, we stained BUD-ELMs with SpyCatcher-GFP and imaged them using confocal microscopy. At the half a millimeter length scale (Fig. 2A – left), the BUD protein (cyan) and cells (yellow) appear distributed throughout the entire material. At the micron length scale, *C. crescentus* cells in the material display a layer of BUD protein (Fig. 2A – right) like planktonic cells (Fig. 1D). However, at the tens of micron length scale (Fig. 2A – middle), we unexpectedly observed a BUD protein-containing secreted matrix (blue) that was locally inhomogeneous and was surrounded by *C. crescentus* cells (yellow) on all sides (Fig. S4). To probe the matrix composition, we also stained the BUD-ELM with Congo Red and 3,3’-dioctadecyloxacarbocyanine perchlorate (DiO) (Fig. 2B), which are known to bind amyloid proteins^28^ and lipids^29^, respectively. Congo Red staining (Fig. 2B – *Proteins*) was orthogonal to the cell channel and analogous to the SpyCatcher-GFP staining, confirming that the matrix is made of proteins. In contrast, DiO (Fig. 2B – *Lipids*), did not stain cell-free regions. Analysis of the cell-free and stained areas (Fig. 2C) confirmed that protein staining had a higher overlap with cell-excluded matrix regions compared to lipid staining. Thus, the BUD-ELM strain produces a secreted proteinaceous matrix containing the BUD protein that mediates BUD-ELM structure at the tens of micron length scale.

**Fig. 2.**
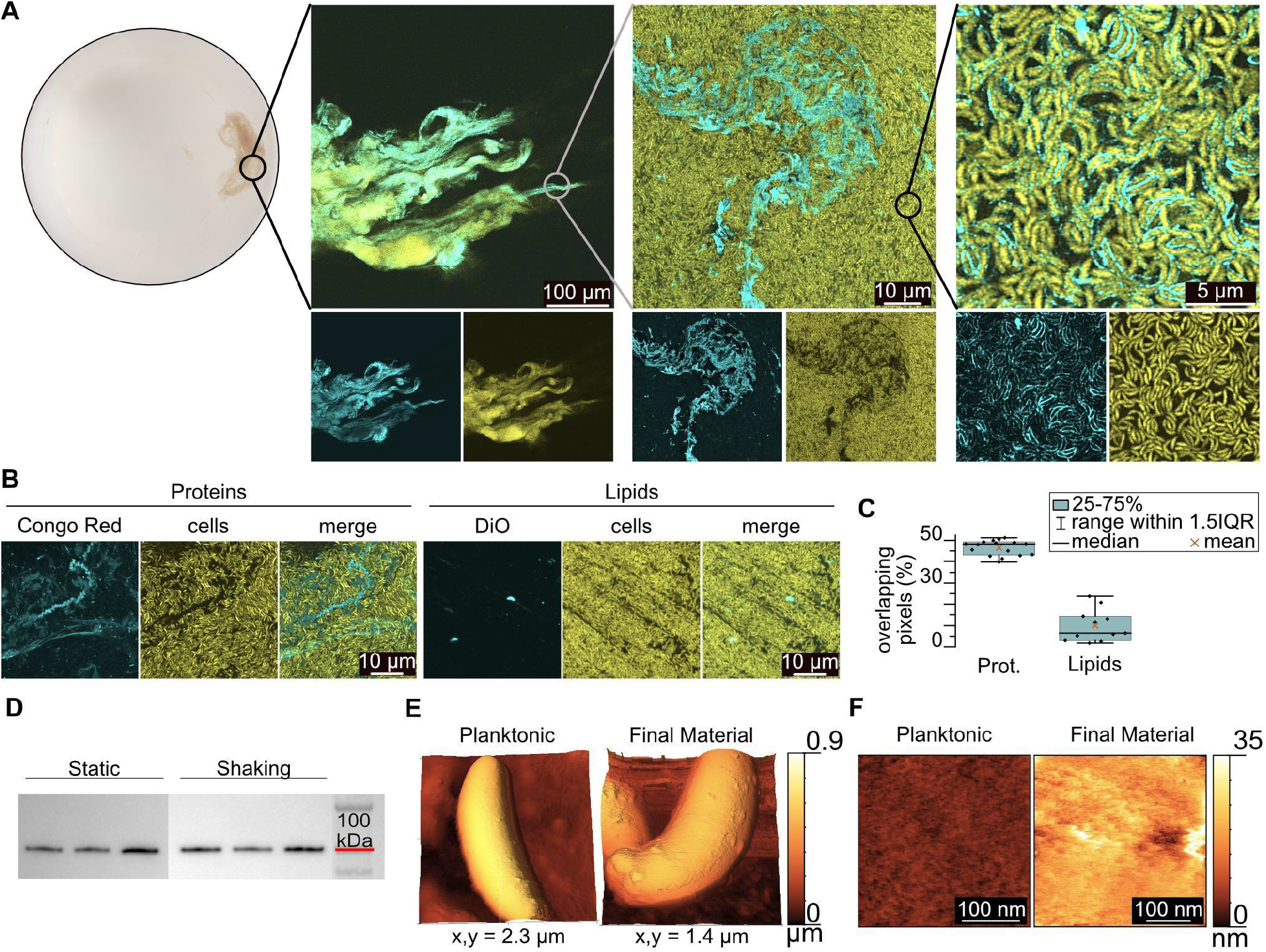
BUD-ELMs contain a *de novo* protein matrix and display hierarchical structure. **(A)** Confocal microscopy of ELMs stained with SpyCatcher-GFP at increasing magnifications, showing a hierarchical structure. Bottom images show individual fluorescent channels: GFP (matrix) on the left and mKate2 (cells) on the right. (**B**) Confocal microscopy of BUD-ELMs stained with Congo Red (left panel) and DiO (right panel), highlighting the proteinaceous nature of the BUD protein matrix. (**C**) Percentage of overlapping pixels between cell-free and stained regions, confirming the absence of lipids in the BUD protein matrix. Error bars represent standard error of 16 images for the protein and 11 for the lipid staining. (**D**) Immunoblot of BUD protein in growth media in static (left) and shaking (right) cultures. (**E**) AFM images of single cells at early (left) and late (right) stages of BUD-ELM formation, showing a difference in surface morphology. (**F**) High-resolution AFM images of single-cell surfaces at early (left) and late (right) stages of BUD-ELM formation, showing differences in surface layer thickness.

To understand how the BUD protein could be both a surface-displayed and secreted matrix, we imaged single cells through AFM at early stages of BUD-ELM formation, when cells are mostly in the planktonic state, and at later stages when the material is fully assembled. At the early stage (Fig. 2E – left), the cell surface appeared uniform, but after the BUD-ELM had formed, cells showed large protuberances (Fig. 2E – right). Additionally, the surface layer depth of early-stage BUD-ELM cells is ∼10 nm (Fig. 2F – left), compared to the ∼35 nm layer of late-stage cells (Fig. 2F – right), indicating that the protein layer thickens over time. We found secreted BUD protein in the medium under both static and shaking conditions (Fig. 2D), indicating that some BUD protein is released into the medium independent of shaking. Moreover, a strain that only secretes, but does not display the BUD protein, created material with a much lower cell content than the BUD-ELM (Fig. S5). These results suggest that the BUD protein accumulates on the cell surface, some of which detaches and forms a secreted matrix that subsequently binds *C. crescentus* cells. We propose that the ability of cells to simultaneously self-interact and adhere to the matrix plays a pivotal role in creating an emergent structure that is cell-rich and macroscopic.

We next sought to understand how this material assembles by imaging BUD-ELM cultures at various times during their growth (Fig. 3A). Shaken cultures grew planktonically for ∼ 12 h (Fig. 3A – left) before a thin pellicle appeared at the air-water interface (Fig. 3A – middle). AFM images of the pellicle depicted a central, cell-dense region (Fig. 3B – left) and a peripheral region of a few cells attached to a ∼6 nm thick membrane (Fig. 3B – middle and right), suggesting the BUD protein forms a protein membrane to which cells adhere. The pellicle increased in density and opacity, becoming more compact. After ∼24 h total culturing time, the pellicle desorbed from the air-water interface and sank as the final material (Fig. 3A – right). Disrupting the hydrophobicity of the air/water interface by the addition of surfactant prevented pellicle and material formation (Fig. S6). Similarly, neither a pellicle nor material formed under static growth conditions. However, when static cultures were shaken, a pellicle formed (Fig. S7). Together, these experiments demonstrate that BUD-ELMs are formed through a multi-step process and establish hydrophobicity of the air/water interface and shaking as critical conditions for assembly of BUD-ELMs.

**Fig. 3.**
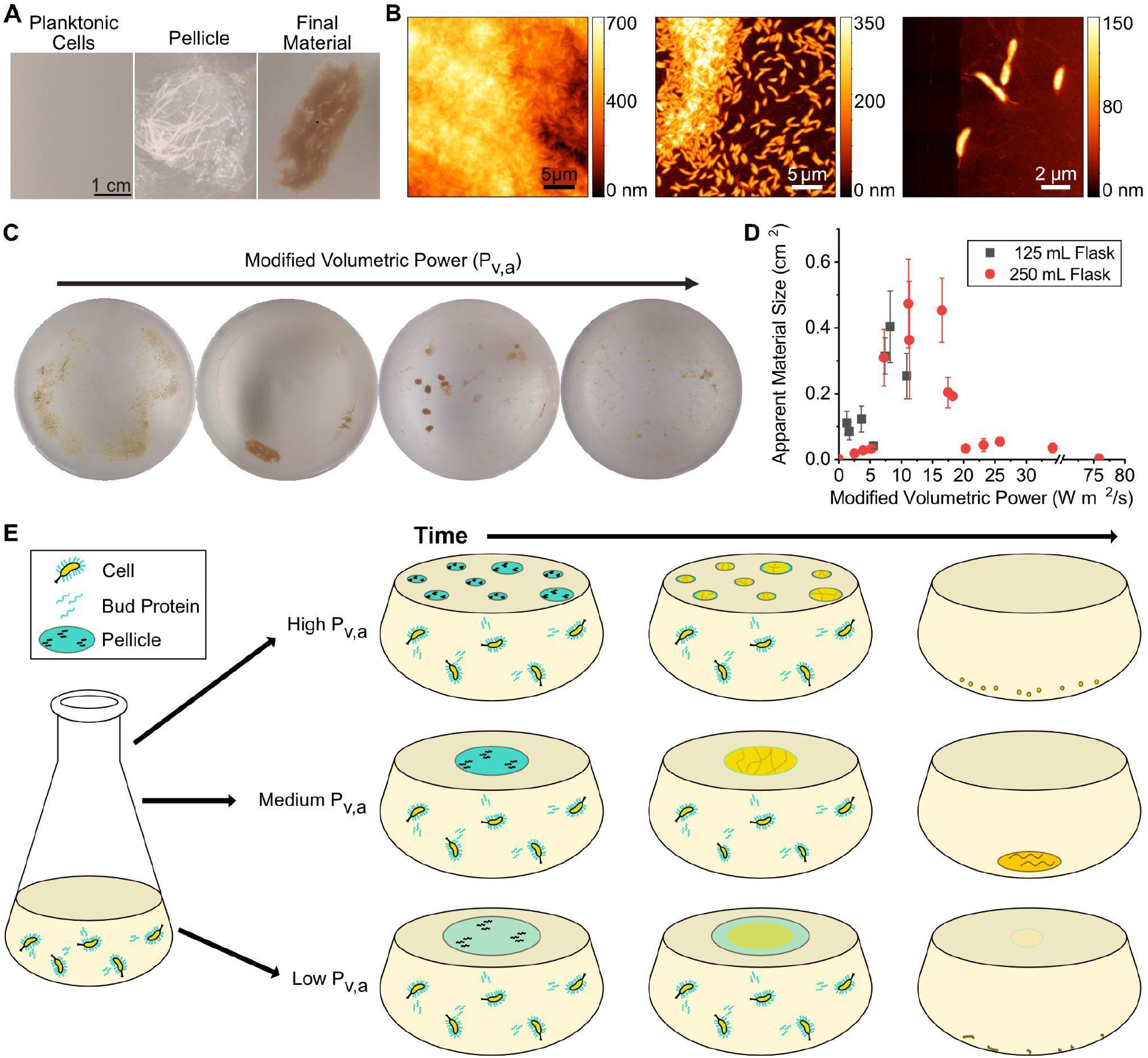
BUD-ELMs are formed through a shaking-dependent, multi-step process. (**A**) Optical images of representative BUD-ELM strain culture during material, showing BUD-ELMs are formed through a multi-step process. (**B**) AFM images of pellicle structure, showing the pellicle contains both a central region containing several layers of densely packed cells (left), and a peripheral region containing sparse cells connected by a thin membrane (center and right). (**C**) Representative optical images of BUD-ELMs grown under different modified volumetric power values. Altering the modified volumetric power changes the morphology and size of the BUD-ELMs. (**D**) Correlation between modified volumetric power and the apparent surface area of the BUD-ELMs for 125- and 250-mL shake flasks. Error bars represent standard error of at least three samples. (**E**) Proposed mechanism for BUD-ELM formation.

### BUD-ELMs are formed through a multi-step process that depends on physical parameters

To understand how physical parameters affect BUD-ELM assembly, we grew cultures under different conditions and measured the size of the resulting materials. The size of BUD-ELMs depended non-monotonically on the shaking speed, volume, and flask diameter (Fig. 3C). We then used these parameters to calculate two quantities: the volumetric power input, P_V_, describing the energy provided to the flask by shaking per unit volume and the volumetric mass transfer coefficient, k_La_, representing the transfer of oxygen into the medium relative to the area of the air-water interface. We found that neither parameter showed a consistent relationship with the size of BUD-ELMs per flask (Fig. S8). Instead, we found empirically that the product of P_V_, k_La_, and the fifth power of the flask diameter, which we refer to as the modified volumetric power parameter, P_V,A_, related the culture conditions to the material size (Fig. 3D). With these data, we propose a model for BUD-ELM assembly (Fig. 3E). During culturing, the BUD protein accumulates in solution and on the surface of *C. crescentus.* With shaking, the BUD protein adsorbs to the air-water interface to form a protein-rich membrane of increasing thickness. BUD protein-displaying cells adhere to the membrane, increasing its density to form a pellicle. Hydrodynamic forces from shaking cause the pellicle to collapse on itself, until the material sinks to the bottom of the flask. At lower P_V,A_ values, the weaker hydrodynamic forces do not collapse the pellicle, and the pellicle fragments into smaller materials. At intermediate P_V,A_ values, stronger hydrodynamic forces collapse the pellicle into a single, large BUD-ELM. At higher P_V,A_ values, shear forces prevent the assembly of larger pellicles, leading to smaller pellicles and final materials. This empirical model provides a basis for the future development of mechanistic models describing BUD-ELM assembly.

### BUD-ELMs are self-regenerating, multifunctional materials whose mechanical properties can be tuned genetically

Since the matrix plays an important role in determining the mechanics of biomaterials, and the matrix of BUD-ELMs is mostly constituted by the BUD proteins, we hypothesized that genetic manipulations of the BUD protein will have a significant effect on the mechanical properties of BUD-ELMs. To test this, we interrogated the effects of removing the flexible biopolymer domain (strain *ΔELP_60_ΔSpyTag,* Fig. 4A – top) and the anchoring domain (strain Δ*rsaA_1-250_*, Fig. S5A) from the original BUD-ELM strain. We observed that both the *ΔELP_60_ΔSpyTag* (Fig. S5B – right) and Δ*rsaA_1-250_* strain (Fig. 4A – bottom) form BUD-ELMs, that are morphologically distinguishable from the original BUD-ELM material in the flask after growth. However, despite these apparent differences, rheological measurements (Fig. S9) confirm that all three BUD-ELMs are viscoelastic solids. Frequency sweep curves (Fig. S10) show large and significant differences in the storage modulus (G′) and the loss modulus (G″) among the three BUD-ELMs throughout the tested range of angular frequency. For a central value of angular frequency of 10 rad/s (Fig. 4B), the storage modulus of *ΔELP_60_ΔSpyTag* BUD_ELMs is ∼340% of the original BUD-ELM, whereas the loss modulus increased by ∼ 300%. Conversely, the Δ*rsaA_1-250_* BUD-ELMs show a lower G’ and G’’, respectively 69% and 84% less than the original BUD-ELM. We speculate that the increased stiffness of the *ΔELP_60_ΔSpyTag* BUD-ELMs reflects the removal of a long flexible linker, the ELP, from the cohesive BUD protein forming this cellular material. On the other end, we suggest Δ*rsaA_1-250_* BUD-ELMs are less stiff due to the lack of crosslinking among cells and between the matrix and the cells. Overall, these results demonstrate that we can control BUD-ELMs mechanical properties over a 16-fold range through genetic modification of the matrix-forming BUD-protein.

**Fig. 4.**
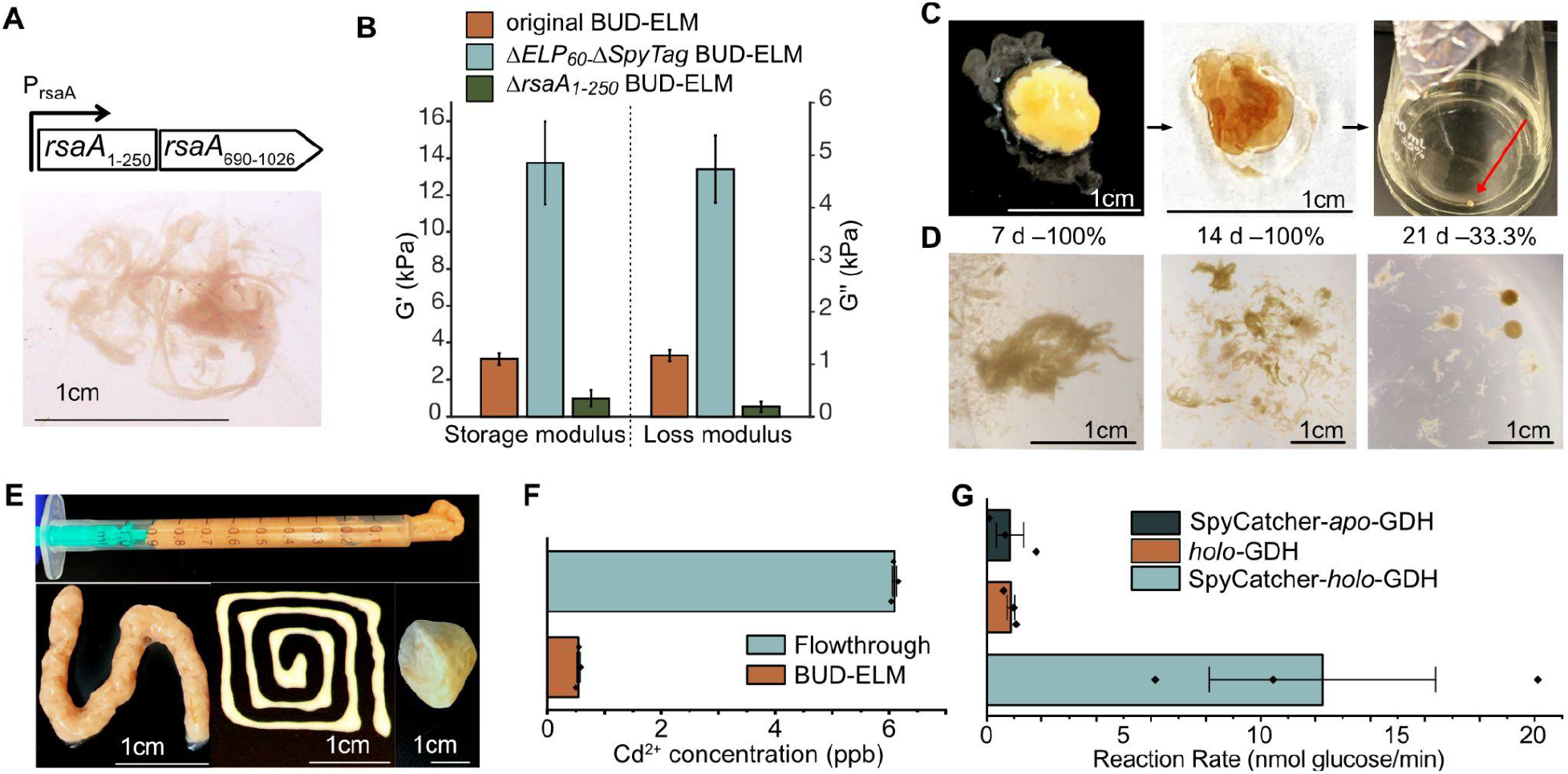
BUD-ELMs are self-regenerating, processible, and functional materials. (**A**) Genetic constructs replacing the native *rsaA* gene in the *ΔELP_60_ΔSpyTag* (left), and image of corresponding BUD-ELM (right). (**B**) Storage (G’) and Loss (G’’) modulus of original, *ΔELP_60_ΔSpyTag* and Δ*rsaA*_1-250_ BUD-ELMs at an angular frequency of 10 rad s^-1^. Error bars represent 95% confidence intervals of at least five samples. (**C**) Representative example of BUD-ELMs reseeding, showing extraction from liquid culture (left), desiccated (middle) and inoculation into fresh medium (right). (**D**) Representative example of BUD-ELMs grown from desiccated material after 7 (left), 14 (middle) or 21 (right) days. The percentage of successful BUD-ELM regeneration was 100%, 100% and 33.3% respectively. Percentages are calculated from at least nine samples. (**E**) BUD-ELMs collected into a syringe (top) for extrusion using different-sized nozzles, (bottom-left and bottom-middle), showing their ability to be reshaped. BUD-ELMs mixed with glass powder to form a firm paste that hardens when dehydrated (bottom-right), showing its potential as cement-like agent. (**F**) Graph showing the final Cd^2+^ solution concentration after 6 ppb Cd^2+^ solution was incubated with (BUD-ELM) or without (flowthrough) the *ΔSpyTag* BUD-ELMs. Error bars represent standard error of three samples. (**G**) Graph showing the rate of glucose oxidation for BUD-ELMs that were incubated with SpyCatcher-*holo*-GDH, *holo*-GDH, or SpyCatcher-*apo*-GDH. Error bars represent standard error of three samples.

ELMs must be able to be processed and stored without losing their ability to regrow. We dried BUD-ELMs (Fig. 4C – left and middle) and re-inoculated fragments of them into fresh medium (Fig. 4C – right). BUD-ELM fragments dried for 7, 14, or 21 days regenerated to form additional BUD-ELMs (Fig. 4D). Whereas BUD-ELMs re-grew 100% of the time after 7 or 14 days of drying, BUD-ELMs desiccated for 21 days regenerated in 33% of cases. Additionally, BUD-ELMs collected from multiple cultures formed a cohesive paste (Fig. 4E – top) that was extrudable through syringes with different diameters (Fig. 4E – bottom, left and middle). When mixed with glass powder, BUD-ELMs created a firmer paste that hardened into a solid composite (Fig. 4E – bottom, right). These results indicate that BUD-ELMs can regenerate after drying, can be reshaped, and can be processed into composite materials.

Lastly, we probed the ability of BUD-ELMs to behave as functional materials. Self-regenerating materials that remove heavy metals from water could help address the growing prevalence of heavy metal contamination. Since many forms of biomass non-specifically absorb heavy metals, we hypothesized that the BUD-ELM could remove Cd^2+^ from solution. When 0.013±0.007g of *ΔSpyTag* BUD-ELM was incubated for 90 min with a CdCl_2_ solution of 6 ppb– 1 ppb above the Environmental Protection Agency (EPA) limit–90 ±5 % of cadmium was removed (Fig. 4F). Next, we functionalized the BUD-ELM matrix to allow it to perform biological catalysis. We fused the oxidoreductase PQQ-glucose dehydrogenase (GDH), which couples oxidation of glucose to reduction of a soluble electron carrier^30^, to SpyCatcher. Cell lysates containing over-expressed *apo* SpyCatcher-GDH or GDH were reconstituted by adding the cofactor PQQ (pyrroloquinoline quinone) to obtain the *holo* forms of the enzyme. After confirming the activity of *holo* GDH in both cases, we observed that only BUD-ELMs incubated with SpyCatcher-*holo*-GDH enzymatically reduced an electron carrier (Fig. 4G and Fig. S12A). This demonstrates that BUD-ELM can be functionalized directly from complex mixtures to act as catalysts. Together, these results show BUD-ELMs can serve as versatile functional materials.

In this work, the serendipitous creation of macroscopic ELMs allowed us to identify a new design principle - that macroscale BUD-ELMs are associated with a secreted extracellular matrix. We suggest that cell-matrix interactions may be essential for BUD-ELMs to reach a macroscopic size. This idea is supported by previous literature that shows that neither cell-cell adhesion alone^31^, nor sparse cell-matrix interactions in the absence of additional forces^19^ lead to microscopic cell aggregates. Additionally, we have demonstrated that nucleation of a pellicle at the liquid-air interface and hydrodynamically-driven coalescence and collapse of the pellicle are required to form macroscopic ELMs. Since pellicle formation is also a key step in nanocellulose-based living materials^18^, we suggest that the use of the air-water interface to locally concentrate and order hydrophobic biomolecules into a matrix may represent a general assembly principle for macroscopic ELMs. The new tools and *C. crescentus* platform developed here will permit systematic exploration of design and assembly rules for programming the growth of centimeter-scale structures using living cells as building blocks.

By creating BUD-ELMs with a *de novo*, modular protein matrix, this work greatly expands the ability to tailor macroscopic ELMs for specific applications. Existing examples of macroscopic, bottom-up ELMs have extracellular matrices predominantly composed of polysaccharides, allowing little control over composition or mechanics^18^. Here we show genetic changes to a single engineered protein can yield dramatic changes in the ELM’s composition and mechanical properties. The modularity of the BUD protein and the ease of engineering protein biopolymers offer much greater opportunities for introducing desirable properties into the matrix^11^. Known polypeptides and proteins can exhibit desirable optical, electrical, mechanical, thermal, transport, and catalytic properties^32^. We envision specific matrix properties that can be combined synergistically with existing cellular functions such as sensing, biomolecule production, and information processing. Thus, this work multiplies the opportunities to program ELMs tailored for applications in human health, energy, and the environment.

## Acknowledgements

We thank Marimikel Charrier, Maria Orozco Hidalgo, and Dr. Vera Troselj for helpful conversations.

## Author contributions

Conceptualization: SM, RFT, DL, CAF, KRR, PDA

Methodology: SM, RFT, RC, SS, JS, DL, CAF

Investigation: SM, RFT, RC, SS, JS, DL

Visualization: SM, RFT, RC, DL

Funding acquisition: CAF, KRR, PDA

Project administration: CAF

Supervision: SM, CAF

Writing – original draft: SM, CAF, RFT

Writing – review & editing: SM, CAF, RFT, RC, SS, JS, DL, KRR, PDA

## Data and materials availability

All strains and plasmids can be supplied upon reasonable request.

## Supplementary Materials

Materials and Methods

Figs. S1 to S12

Tables S1 to S3

## Materials and Methods

### Construction of C. crescentus strains

All *C. crescentus* strains used in this work are listed in Table S1.

To create the *C. crescentus* strains, we cloned integration plasmids designed to incorporate synthetic genes into the *rsaA* locus using homologous recombination. For the BUD-ELM strain (RCC002), we created integration plasmid pSMCAF008 by inserting a target sequence into the multicloning site of the backbone plasmid pNPTS138 (GenBank: MK533795.1) using restriction enzymes (ApaI upstream and NheI downstream). The target sequence (ordered from GenScript, sequence below) contained the DNA sequence encoding the *ELP_60_-SpyTag* flanked by 800bp of homology regions up- and downstream of the native *rsaA* central domain (*rsaA* 750-2073). These flanking regions allow the synthetic gene to be properly positioned within the *C. crescentus* genome. The *ΔELP_60_*Δ*SpyTag* and Δ*SpyTag* strains (RCC004 and RCC005) were generated with integration plasmids pSMCAF017 and pSMCAF018. These plasmids were cloned by PCR amplification of pSMCAF008 using the primers listed below and Golden Gate assembly.

The wild type strain of *C. crescentus* (MFm126) was transformed with pSMCAF008, pSMCAF017 and pSMCAF018, to generate RCC002, RCC003, and RCC004 strains, respectively, using a two-step recombination technique^20^. The two-step recombination technique with sucrose counterselection is as follows: the pNPTS138 plasmids were electroporated into *E. coli* WM3064 cells and subsequently conjugated overnight into *C. crescentus* NA1000 *ΔsapA*::Pxyl-mkate2 (MFm126) on a PYE agar plate containing 300n µM DAP. The culture was then plated on PYE with kanamycin to select for integration of the plasmid and removal of *E. coli* cells. Successful integrants were incubated in liquid PYE media overnight and plated on PYE supplemented with 3% w/v sucrose to select for excision of the plasmid and *sacB* gene, leaving the target sequence in the genome. Integration of the sequences was confirmed by colony PCR (with primers SMCAF078 and SMCAF079) using a Touchdown thermocycling protocol with an annealing temperature ranging from 72-62°C, decreasing 1°C per cycle. The PCR amplicons have been later sequenced (with primers SMCAF093 and SMCAF095);

Primers used to verify the synthetic strains: SMCAF078:

CTTAGTCTAGCGATCCTCGCCTAG

SMCAF079: ATCGCTGCTCCCATGCGC

SMCAF093: GTCCTTGTAGTCACCCGAG

SMCAF095: AGTTGCAGAGCCGTGAAG

Primers used to assemble pSMCAF017:

SMCAF142: TGGGTCTCAAGGGCGGTTCGGGAGGAGGC

SMCAF143: TGGGTCTCAGCAATCCAAACGAGAGTCTAATAGAATGAGGTC

SMCAF144: TGGGTCTCCTTGCAACTGGTCTATTTTCCTCTTTTG

SMCAF145: TAGGTCTCCCCCTTGTCATCGTCGTCCTTG

Primers used to assemble pSMCAF018:

SMCAF168: CAGGTCTCTTGTCGCACCTGATTGCCCGA

SMCAF169: CAGGTCTCTGCTGGGACACCACCGCCAGG

SMCAF170: CTGGTCTCCCAGCTGACCCGGCCTTCGGC

SMCAF171: CTGGTCTCTGACAATCTATCGATTGTATGGGAAGCCCG

GenScript target sequence:

CCAATGATCGTAATACGACTCACTAGTGGGGCCCGCGCCACTCGGTCGCAGGGGGTGTGGGATTTTTTTTGGGAGACAATCCTCATGGCCTATACGACGGCCCAGTTGGTGACTGCGTACACCAACGCCAACCTCGGCAAGGCGCCTGACGCCGCCACCACGCTGACGCTCGACGCGTACGCGACTCAAACCCAGACGGGCGGCCTCTCGGACGCCGCTGCGCTGACCAACACCCTGAAGCTGGTCAACAGCACGACGGCTGTTGCCATCCAGACCTACCAGTTCTTCACCGGCGTTGCCCCGTCGGCCGCTGGTCTGGACTTCCTGGTCGACTCGACCACCAACACCAACGACCTGAACGACGCGTACTACTCGAAGTTCGCTCAGGAAAACCGCTTCATCAACTTCTCGATCAACCTGGCCACGGGCGCCGGCGCCGGCGCGACGGCTTTCGCCGCCGCCTACACGGGCGTTTCGTACGCCCAGACGGTCGCCACCGCCTATGACAAGATCATCGGCAACGCCGTCGCGACCGCCGCTGGCGTCGACGTCGCGGCCGCCGTGGCTTTCCTGAGCCGCCAGGCCAACATCGACTACCTGACCGCCTTCGTGCGCGCCAACACGCCGTTCACGGCCGCTGCCGACATCGATCTGGCCGTCAAGGCCGCCCTGATCGGCACCATCCTGAACGCCGCCACGGTGTCGGGCATCGGTGGTTACGCGACCGCCACGGCCGCGATGATCAACGACCTGTCGGACGGCGCCCTGTCGACCGACAACGCGGCTGGCGTGAACCTGTTCACCGCCTATCCGTCGTCGGGCGTGTCGGGTTCGGGCGGTTCGGGAGGAGGCTCGGGTGACTACAAGGACGACGATGACAAGGGAGTTGGCGTCCCAGGAGTTGGAGTCCCAGGAGGGGGCGTTCCGGGCGCAGGAGTTCCTGGAGTAGGAGTTCCAGGAGTGGGCGTGCCAGGGGTGGGCGTCCCAGGTGGGGGAGTTCCCGGAGCAGGTGTGCCTGGGGGCGGCGTGCCTGGAGTCGGAGTTCCGGGGGTGGGTGTACCGGGTGGAGGCGTACCAGGCGCGGGAGTGCCGGGCGTGGGCGTGCCAGGCGTCGGTGTACCCGGCGTTGGTGTTCCGGGCGGAGGTGTCCCCGGAGCTGGGGTTCCCGGTGGGGGTGTACCGGGCGTCGGGGTTCCCGGTGTGGGTGTCCCAGGTGGCGGCGTTCCCGGGGCGGGCGTA

CCTGGAGTGGGTGTGCCAGGAGTCGGCGTCCCAGGAGTCGGCGTACCAGGAGGTGGTGTTCCCGGGGCCGGAGTTCCCGGCGGAGGAGTTCCCGGCGTCGGCGTCCCTGGGGTCGGCGTCCCGGGAGGTGGAGTACCCGGAGCAGGAGTGCCGGGAGTCGGTGTACCTGGTGTCGGTGTCCCTGGTGTAGGTGTCCCGGGTGGTGGGGTGCCAGGTGCTGGCGTACCTGGGGGGGGGGTTCCTGGCGTAGGCGTTCCGGGGGTGGGCGTTCCGGGCGGCGGGGTGCCGGGAGCAGGTGTCCCCGGCGTTGGTGTACCGGGGGTTGGTGTCCCAGGCGTAGGTGTGCCCGGTGGAGGGGTGCCGGGAGCTGGAGTGCCTGGAGGGGGTGTACCAGGGGTCGGTGTTCCCGGTGTAGGAGTACCGGGGGGCGGAGTCCCAGGAGCCGGCGTGCCGGGTGTTGGAGTCCCGGGAGTCGGAGTCCCTGGGGTAGGCGTTCCAGGGGGAGGGGTCCCCGGTGCAGGGGTTCCTGGCGGTGGTGTCCCAGGCGGTTCGGGAGGAGGCTCGGGTGCGCATATCGTAATGGTCGATGCATACAAGCCCACGAAAGGAGGTTCAGGCGGCGGAAGCGGTGGTGGAAGCGGAGGTGGGTCAGGCGGAGGCTCAGGGGGAGGTTCGGGTGGCGGTTCGGGAGGAGGCTCGGGTGCTGACCCGGCCTTCGGCGGCTTCGAAACCCTCCGCGTCGCTGGCGCGGCGGCTCAAGGCTCGCACAACGCCAACGGCTTCACGGCTCTGCAACTGGGCGCGACGGCGGGTGCGACGACCTTCACCAACGTTGCGGTGAATGTCGGCCTGACCGTTCTGGCGGCTCCGACCGGTACGACGACCGTGACCCTGGCCAACGCCACGGGCACCTCGGACGTGTTCAACCTGACCCTGTCGTCCTCGGCCGCTCTGGCCGCTGGTACGGTTGCGCTGGCTGGCGTCGAGACGGTGAACATCGCCGCCACCGACACCAACACGACCGCTCACGTCGACACGCTGACGCTGCAAGCCACCTCGGCCAAGTCGATCGTGGTGACGGGCAACGCCGGTCTGAACCTGACCAACACCGGCAACACGGCTGTCACCAGCTTCGACGCCAGCGCCGTCACCGGCACGGGCTCGGCTGTGACCTTCGTGTCGGCCAACACCACGGTGGGTGAAGTCGTCACGATCCGCGGCGGCGCTGGCGCCGACTCGCTGACCGGTTCGGCCACCGCCAATGACACCATCATCGGTGGCGCTGGCGCTGAC

ACCCTGGTCTACACCGGCGGTACGGACACCTTCACGGGTGGCACGGGCGCGGATATCTTCGATATCAACGCTATCGGCACCTCGACCGCTTTCGTGACGATCACCGACGCCGCTGTCGGCGACAAGCTCGACCTCGTCGGCATCTCGACGAACGGCGCTATCGCTGACGGCGCCTTCGGCGCTGCGGTCACCCTGGGCGCTGCTGCGACGCTAGCTGACTGGGAAAACCCTGGCGTTAATCGGAAAGAACATGTGAGCAAAAGGCCAGCAAAAGGCCAGGAACCGTAAAAAGGCCGCGTTGCTGGCGTTTTTCCATAGGCTCCGCCCCCCTGACGAGCATCACAAAAATCGACGCTCAAGTCAGAGGTGGCGAAACCCGACAGGACTATAAAGATACCAGGCGTTTCCCCCTGGAAGCTCCCTCGTGCGCTCTCCTGTTCCGACCCTGCCGCTTACCGGATACCTGTCCGCCTTTCTCCCTTCGGGAAGCGTGGCGCTTTCTCATAGCTCACGCTGTAGGTATCTCAGTTCGGTGTAGGTCGTTCGCTCCAAGCTGGGCTGTGTGCACGAACCCCCCGTTCAGCCCGACCGCTGCGCCTTATCCGGTAACTATCGTCTTGAGTCCAACCCGGTAAGACACGACTTATCGCCACTGGCAGCAGCCACTGGTAACAGGATTAGCAGAGCGAGGTATGTAGGCGGTGCTACAGAGTTCTTGAAGTGGTGGCCTAACTACGGCTACACTAGAAGAACAGTATTTGGTATCTGCGCTCTGCTGAAGCCAGTTACCTTCGGAAAAAGAGTTGGTAGCTCTTGATCCGGCAAACAAACCACCGCTGGTAGCGGTGGTTTTTTTGTTTGCAAGCAGCAGATTACGCGCAGAAAAAAAGGATCTCAAGAAGATCCTTTGATCTTTTCTACGGGGTCTGACGCTCAGTGGAACGAAAACTCACGTTAAGGGATTTTGGTCATGAGATTATCAAAAAGGATCTTCACCTAGATCCTTTTAAATTAAAAATGAAGTTTTAAATCAATCTAAAGTATATATGAGTAAACTTGGTCTGACAGTTACCAATGCTTAATCAGTGAGGCACCTATCTCAGCGATCTGTCTATTTCGTTCATCCATAGTTGCCTGACTCCCCGTCGTGTAGATAACTACGATACGGGAGGGCTTACCATCTGGCCCCAGTGCTGCAATGATACCGCGAGACCCACGCTCACCGGCTCCAGATTTATCAGCAATAAACCAGCCAGCCGGAAGGGCCGAGCGCAGAAGTGGTCCTGCAACTTTATCCGCCTCCA

TCCAGTCTATTAATTGTTGCCGGGAAGCTAGAGTAAGTAGTTCGCCAGTTAATAGTTTGCGCAACGTTGTTGCCATTGCTACAGGCATCGTGGTGTCACGCTCGTCGTTTGGTATGGCTTCATTCAGCTCCGGTTCCCAACGATCAAGGCGAGTTACATGATCCCCCATGTTGTGCAAAAAAGCGGTTAGCTCCTTCGGTCCTCCGATCGTTGTCAGAAGTAAGTTGGCCGCAGTGTTATCACTCATGGTTATGGCAGCACTGCATAATTCTCTTACTGTCATGCCATCCGTAAGATGCTTTTCTGTGACTGGTGAGTACTCAACCAAGTCATTCTGAGAATAGTGTATGCGGCGACCGAGTTGCTCTTGCCCGGCGTCAATACGGGATAATACCGCGCCACATAGCAGAACTTTAAAAGTGCTCATCATTGGAAAACGTTCTTCGGGGCGAAAACTCTCAAGGATCTTACCGCTGTTGAGATCCAGTTCGATGTAACCCACTCGTGCACCCAACTGATCTTCAGCATCTTTTACTTTCACCAGCGTTTCTGGGTGAGCAAAAACAGGAAGGCAAAATGCCGCAAAAAAGGGAATAAGGGCGACACGGAAATGTTGAATACTCATACTCTTCCTTTTTCAATATTATTGAAGCATTTATCAGGGTTATTGTCTCATGAGCGGATACATATTTGAATGTATTTAGAAAAATAAACAAATAGGGGTTCCGCGCACATTTCCCCGAAAAGTGCCACCTGACGTC

### Construction of Escherichia coli strains for protein overexpression

To create *E. coli* strains that overexpressed GFP, GFP-Spycatcher, GDH, and GDH-SpyCatcher, we first constructed plasmids coding for these proteins under control of an arabinose inducible promoter. Plasmids pSMCAF015 and pSMCAF016 were assembled from previously constructed pBAD-RFP^20^ and pBAD-SpyCatcher-RFP^20^ plasmids respectively, by substituting the mRFP sequence with the GFP sequence. Similarly, plasmids pSMCAF032 and pSMCAF029 were assembled from backbones pBAD-RFP^20^ and pBAD-SpyCatcher-RFP^20^ respectively, by substituting the mRFP sequence with the GDH sequence. These plasmids were transformed into chemically competent BL21(DE3) cells (New England Biolabs) and single transformants selected for using ampicillin resistance.

GFP sequence:

ATGCGTAAAGGCGAAGAGCTGTTCACTGGTGTCGTCCCTATTCTGGTGGAACTGGATGGTGATGTCAACGGTCATAAGTTTTCCGTGCGTGGCGAGGGTGAAGGTGACGCAACTAATGGTAAACTGACGCTGAAGTTCATCTGTACTACTGGTAAACTGCCGGTTCCTTGGCCGACTCTGGTAACGACGCTGACTTATGGTGTTCAGTGCTTTGCTCGTTATCCGGACCATATGAAGCAGCATGACTTCTTCAAGTCCGCCATGCCGGAAGGCTATGTGCAGGAACGCACGATTTCCTTTAAGGATGACGGCACGTACAAAACGCGTGCGGAAGTGAAATTTGAAGGCGATACCCTGGTAAACCGCATTGAGCTGAAAGGCATTGACTTTAAAGAAGACGGCAATATCCTGGGCCATAAGCTGGAATACAATTTTAACAGCCACAATGTTTACATCACCGCCGATAAACAAAAAAATGGCATTAAAGCGAATTTTAAAATTCGCCACAACGTGGAGGATGGCAGCGTGCAGCTGGCTGATCACTACCAGCAAAACACTCCAATCGGTGATGGTCCTGTTCTGCTGCCAGACAATCACTATCTGAGCACGCAAAGCGTTCTGTCTAAAGATCCGAACGAGAAACGCGATCATATGGTTCTGCTGGAGTTCGTAACCGCAGCGGGCATCACGCATGGTATGGATGAACTGTACAAATAA

GDH sequence:

GACGTTCCGCTGACCCCGAGCCAGTTTGCGAAAGCGAAAAGCGAGAACTTCGACAAAAAAGTCATCCTGAGCAACCTGAATAAACCGCACGCTCTGCTGTGGGGTCCGGATAATCAGATTTGGCTGACCGAACGCGCAACCGGTAAAATTCTGCGCGTTAACCCGGAAAGCGGCAGCGTTAAAACCGTCTTTCAGGTTCCGGAAATCGTTAACGACGCAGACGG

TCAAAACGGTCTGCTGGGTTTTGCGTTTCATCCGGACTTCAAAAACAACCCGTACATCTACATCAGCGGCACCTTCAAAAACCCGAAAAGTACCGACAAAGAGCTGCCGAATCAGACCATCATCCGTCGCTATACCTACAACAAAAGCACCGACACCCTGGAAAAACCGGTTGATCTGCTGGCAGGTCTGCCGAGTAGTAAAGATCATCAGAGCGGTCGTCTGGTAATTGGTCCGGACCAGAAAATCTACTATACCATTGGCGATCAGGGCCGTAACCAACTGGCATACCTGTTTCTGCCGAACCAAGCACAACATACCCCGACCCAACAAGAACTGAACGGCAAAGACTACCACACCTACATGGGCAAAGTTCTGCGTCTGAATCTGGACGGTAGCATTCCGAAAGACAACCCGAGCTTCAACGGCGTTGTTAGCCATATCTATACCCTGGGTCACCGTAATCCGCAAGGTCTGGCATTTACCCCGAACGGTAAACTGCTGCAGTCTGAACAGGGTCCGAATTCTGACGACGAAATCAACCTGATCGTTAAAGGCGGCAATTACGGTTGGCCGAACGTTGCAGGCTATAAAGACGATAGCGGCTATGCATACGCGAATTATAGCGCAGCGGCAAACAAAAGCATCAAAGACCTGGCCCAGAACGGTGTTAAAGTTGCAGCAGGCGTTCCGGTTACCAAAGAAAGCGAGTGGACCGGCAAAAACTTTGTTCCGCCGCTGAAAACCCTGTATACCGTCCAGGACACCTACAACTATAACGATCCGACCTGCGGCGAAATGACCTATATTTGCTGGCCGACCGTTGCACCGAGTTCTGCATACGTTTACAAAGGCGGCAAAAAAGCGATCACCGGTTGGGAAAATACCCTGCTGGTTCCGAGTCTGAAACGCGGCGTTATCTTCCGCATCAAACTGGATCCGACCTATAGTACCACCTACGACGATGCCGTTCCGATGTTCAAAAGCAACAACCGTTATCGCGACGTTATTGCAAGTCCGGACGGTAACGTTCTGTACGTTCTGACCGATACCGCAGGTAACGTTCAGAAAGACGACGGTAGCGTTACCAATACCCTGGAAAATCCGGGTAGCCTGATCAAATTCACCTACAAAGCGAAATGA

### Growth conditions for C. crescentus strains and BUD-ELMs

Unless indicated otherwise, BUD-ELMs were grown by inoculating a single colony of *C. crescentus* strains into 80 mL of PYE in a 250 mL glass flask. These cultures were grown at 30°C at 250 rpm, and BUD-ELMs typically formed within ∼24-30 h. To explore the effect of growth parameters on BUD-ELM size, the flask volume, shaking speed, and culture volume were varied from 125-250 mL, 0-250 rpm, and 30-80 mL, respectively. The complete list of conditions tested can be found in Table S3.

To test the ability of BUD-ELMs to re-seed their own growth, a piece of BUD-ELM grown under standard conditions was collected and transferred in a petri dish for 7, 14 or 21 days. The material dried out completely in less than 24 h. A ∼0.3 to 0.5 cm^2^ piece was broken off from the original material and inoculated into 80 mL of PYE. The culture was then incubated at 30°C at 250 rpm. We detected BUD-ELM formation in 48 h for material dried over 7 or 14 d, and in 72 h for material dried over 21 d.

### Expression and purification of SpyCatcher-GFP, and GFP from Escherichia coli

Single colonies of *E. coli* BL21(DE3) harboring pSMCAF015 and pSMCAF016, for overexpression of GFP and SpyCatcher-GFP respectively, were inoculated in 25 mL of RM minimal media with 0.2% w/v glucose and 100 µg/mL ampicillin. After ∼16 h of growth at 37°C and 250 rpm, cells were used to inoculate 0.5 L of RM minimal media with 0.2% v/v glycerol, 100 µg/mL ampicillin and 0.0004% antifoam (Antifoam 204) to a final OD_600_ ∼0.05. The cultures were allowed to grow at 37°C until mid-log phase. Protein production was induced with 0.2% w/v L-arabinose with incubation at 30°C for ∼17 h.

Cells were harvested by centrifugation at 8000xg for 30 min, resuspended in lysis buffer (50 mM Tris pH 8.0, 300 mM NaCl, 5% v/v glycerol, and 10 mM Imidazole) and lysed using using Avestin Emulsiflex C3 Homogenizer. The lysate was centrifuged at 12000xg for 1 h and the supernatant was collected for protein purification. The proteins were purified using Immobilized Metal Affinity Chromatography (IMAC) with a HisTrap FF column and buffers containing 50 mM Tris pH 8.0, 300 mM NaCl, 5% v/v glycerol, and 10-250 mM Imidazole. After protein purity was confirmed by SDS-PAGE, the protein was dialysed into TEV-cleavage buffer (50 mM Tris pH 8.0, 0.5 mM EDTA, 1 mM DTT) and the 6 x His-tag was cleaved using TEV protease by agitation at 4°C for 4 h. The cleaved protein was stored at -80°C in 50 mM NaPO_4_ pH 8.0, 300 mM NaCl and 5% v/v glycerol.

### Imaging and Size Analysis of Macroscopic BUD-ELMs

To image the BUD-ELMs, the bottom of a flask containing the BUD-ELM culture was imaged using a Canon EOS 77D camera. Flasks were positioned within a reflective photobox on a clear plastic surface such that the bottom of the flask was positioned approximately 11.5 cm above the camera lens. Flasks were illuminated from above by white LED strip lights, which were dampened by a thin white polyester fabric. For imaging of the air-water interface, each flask was instead imaged from above via a hole in the light box, and a black backdrop was positioned beneath the flask.

To determine the flat surface area of individual pieces of material under different growth conditions, images of flasks were separated into RGB channels using MATLAB. A subset of the blue channel images was then input into the image classification software ilastik, as a training set for the autocontext workflow. The first stage of training separated images into three different classifications: background, scattered material, and bundled material. Scattered material was defined as overlapping regions of small aggregates not associated with each other, whereas bundled material referred to larger, connected pieces of material. The second stage of training distinguished bundled material from the rest of the image. Both stages of training utilized all 37 features provided within the ilastik workflow. From the results of the training set, the second stage segmentation masks for all blue channel images were calculated, and loaded into MATLAB. From these masks, the flat area of each piece of material was calculated, and the top 5 percentile of size from each image was averaged to yield a representative size measurement. For each image, a conversion rate between pixels and millimeters was determined using the standard flask diameter as a reference point. Size measurements were averaged between samples and plotted with respect to their calculated (d5)(kLa)(P/VL) values.

### Imaging of Microscopic BUD-ELM structure using confocal microscopy

Single colonies of BUD-ELM strain (RCC002) were inoculated in 30 mL PYE with 0.15% D-xylose – to induce the expression of mKate2, in a 125 mL flask and grown for 24 h at 30°C at a shaking speed of 250 rpm. BUD-ELMs of similar dimensions were collected and washed twice with 1 mL of 0.01 M Phosphate-buffered saline (PBS), in a centrifuge tube. They were then incubated in 1 mL of 0.01 M PBS, at 30°C, with the following staining agent: 80 µg of SpyCatcher-GFP or GFP for 1 h, 1% CongoRed (ThermoFisher Scientific – D275) or 100 µg DiO (DiOC18(3) - 3,3’-Dioctadecyloxacarbocyanine Perchlorate) for 20 min. Samples were washed 3 times with 1 mL 0.01 M PBS and then a small amount was placed between a slab of PYE agarose (1.5% w/v) and a glass coverslip-bottomed 50-mm Petri dish with a glass diameter of 30 mm (MatTek Corporation). To acquire the low-magnification images (Fig. 2A – left panel), BUD-ELMs were embedded into 5% w/v agarose and sliced. The slice was placed on a glass coverslip-bottomed 50-mm Petri dish. For imaging, we used the Zeiss LSM800 Airyscan confocal microscope.

### In situ atomic force microscopy (AFM) imaging on living C. crescentus cells

Poly-L-lysine coated silicon substrates were immersed in Falcon™ round-bottom polypropylene culturing tubes containing 3 mL fresh *C. crescentus* cell culture at an OD_600_ of 0.3-0.5. Culture tubes were then centrifuged at 3000×g for 10 min to immobilize the cells onto the silicon substrate. The silicon substrate was washed with 2 mL of sterile PYE to remove loosely-bound cells before being mounted to a metal puck and transferred to the AFM sample stage. *In situ* AFM was performed on an Asylum Cypher AFM using soft tapping mode. A fluid cell and two syringe pumps were assembled to control liquid flow and PYE medium was supplied to maintain cell viability during imaging. The AFM probe consisted of a sharp silicon tip on a silicon nitride cantilever (BioLever mini, BL-AC40TS) with a spring constant of 0.09 N/m. Cells were imaged in native state without fixation. A 100-200 mV amplitude setpoint was used to apply minimum forces (∼0.2 nN) to cells during imaging.

### AFM imaging on pellicle structures

To bind BUD-ELM pellicle to a silicon substrate, a 2 cm^2^ precleaned silicon substrate was dipped into the pellicle forming cell culture with an entry angle of ∼60° perpendicular to the water surface. The silicon substrate was then retrieved, and the pellicle structure was dried under N_2_ atmosphere for 2 h. The dried pellicle structure on the substrate was mounted to a metal puck and transferred to the AFM sample stage. AFM imaging was performed on an Asylum Cypher AFM using soft tapping mode in air. A Tap-150 tip (BudgetSensors) with 5 N/m force constant was used to image the pellicle structure.

### Immunoblot Analysis of BUD proteins

Cultures of *C. crescentus* BUD-ELM strain (RCC002) were cultured in standard or stationary (not shaking) conditions. At 36 h for static cultures and at 24 h (sunken material phase) for shaking cultures, the supernatant of each culture was extracted and loaded onto a TGX Stain-Free™ gel (Bio-rad). After running, the gel was transferred to a 0.2 µm nitrocellulose membrane and blocked for 1 h at room temperature with SuperBlock™ blocking buffer (Thermo Scientific). Membranes were then washed four times in TBST buffer before incubation in a 1:5000 dilution of Monoclonal ANTI-FLAG® antibody (Monoclonal ANTI-FLAG® M2-Peroxidase (HRP) antibody produced in mouse, clone M2, purified immunoglobulin, buffered aqueous glycerol solution) solution for 1 h at room temperature. Membranes were washed an additional four times in TBST buffer before Clarity Max Western ECL Substrate (Bio-rad) was applied and the membrane was imaged for chemiluminescence. Each sample lane represents an independent sample, grown from separate individual colonies.

### Building of Predictive Parameter for Material Size

To describe the effect of shaking on BUD-ELM formation, a model was built based on the volumetric power input of a shake flask. The volumetric power input, defined as the rate of energy transfer into a shake flask per unit volume, was described by Büchs *et al*.^33^ as:

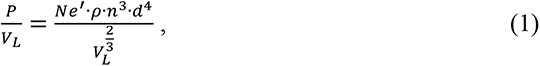

Where *P* is power, *V_L_* is culture volume, *n* is shaking frequency, *d* is the inner flask diameter, and *Ne’* is the modified Newtons number. *Ne’* was written in Büchs *et al.*^33^ as a function of the Reynolds number *Re’* in the following manner:

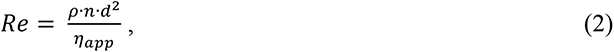

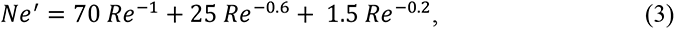

where η_app_ is the apparent dynamic viscosity of the culture. To consider the impact of the air-water interface on BUD-ELM assembly, the volumetric power input was multiplied by the volumetric mass transfer coefficient of oxygen, defined dimensionally by Klöckner and Büchs^34^, as:

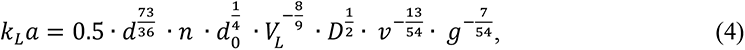

where *k_L_* is the transfer coefficient of oxygen, *a* is the oxygen transfer surface area, *d_0_* is the shaking orbit diameter, *D* is the diffusion coefficient, *v* is the kinematic viscosity, and *g* is the acceleration of gravity. To unify size measurements across different flask sizes, a correction factor of *d*^5^ was applied. This new parameter was dubbed the “modified volumetric power”, defined as:

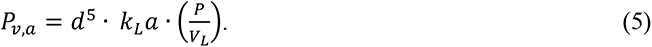

Calculations for *P_v,a_* assumed that the media parameters for viscosity η_app_ and v and the diffusion coefficient *D* were equal to that of water at standard growth temperature of 30°C, which approximates culture conditions at inoculation.

### Parameter definitions and units

*n*– shaking frequency (s^-1^); *Ne*’– modified Newton number (dimensionless); *P* – power input (W); *Re* – Reynold’s number (dimensionless); *V_L_* – culture volume (m^3^); η_app_ – dynamic apparent viscosity (Pa s); ρ – liquid density (kg/m^3^); *d_0_* – orbital shaking diameter (m); *D –* diffusion coefficient (m^2^/s); *v –* kinematic viscosity (m^2^/s); *g* – acceleration of gravity (m/s^2^)

### Rheological measurements

The rheological properties of BUD-ELMs produced from strains RCC002, RCC004 and MFm152 (original BUD-ELM, Δ*ELP_60_ΔStpTag*, and Δ*rsaA_1-250_* strains, respectively) were evaluated on a strain-controlled rheometer (ARIES G2) equipped with an 0.1 rad 8-mm diameter cone plate. BUD-ELMs were grown in standard conditions. An approximate volume of 100-200 µL of BUD-ELMs were collected into a 1.5 mL centrifuge tube and spun for 10 sec at 3200 rcf with a mini centrifuge (VWR®, C0803). This allowed for the material to collect at the bottom as an homogeneous paste. The supernatant was removed and 150uL of fresh PYE were added on top of BUD-ELMs to prevent desiccation. Strain sweep experiments from 0.1 to 100% strain amplitudes were performed at a fixed frequency of 3.14 rad/s. Frequency sweep experiments from 100 to 0.1 rad/s were performed at a 0.35% strain amplitude.

### Biosorption of Cd^2+^ to BUD-ELMs

To measure the Cd^2+^ ability of BUD-ELMs, the *ΔSpyTag* strain was cultured in standard conditions. After growth for between 24 to 48 h, BUD-ELMs were harvested into sterile 2 mL tubes and lyophilized for 5 h. Lyophilized BUD-ELMs were transferred to a metal-free 15 mL tube (VWR® Metal-Free Centrifuge Tubes, Polypropylene, Sterile) and incubated with 7 mL of 6 ppm CdCl_2_ (Sigma Aldrich 202908) in ddH2O for 90 min on an orbital shaker. After incubation, the Cd^2+^ concentration of the supernatant was measured by ICP-MS. Specifically, 5 µL of the supernatant was diluted in 4.995 mL 1% HNO_3_ with 5 µg/mL Indium (In), as standard for data analysis (Perkin Elmer N9303741). This diluted solution was run on a Perkin Elmer Nexion 300 ICP-MS with two isotopic measurements (Cd^2+^ 111, Cd^2+^ 112) and In 115 as the internal standard. Data was analyzed using Syngistix software. Samples were run in biological triplicates and data provided is mean with standard error.

### Functionalization of BUD-ELMs with PQQ-Glucose Dehydrogenase (GDH)

Single colonies of *E. coli* BL21(DE3) harboring pSMCAF032 and pSMCAF029, for overexpression of GDH and SpyCatcher-GDH respectively, were inoculated in 25 mL of Terrific broth (TB) with 100 µg/mL ampicillin. After ∼16 h of growth at 37°C and 250 rpm, cells were used to inoculate 0.5 L of TB with 0.02% antifoam (Antifoam 204) and 100 µg/mL ampicillin to a final OD_600_ ∼0.05. The cultures were allowed to grow at 37°C until mid-log phase. Protein production was induced with 0.2% w/v L-arabinose with incubation at 30°C for ∼17 h.

Cells were harvested by centrifugation at 8000xg for 30 min, resuspended in lysis buffer (0.02 M PBS, 1mM MgCl_2_) and lysed using Avestin Emulsiflex C3 Homogenizer. Lysates were centrifuged at 12000xg for 1 h and the supernatant was collected for BUD-ELM functionalization. The GDH (or SpyCatcher-GDH) in the lysate was reconstituted by adding a final concentration of 3 mM Ca^2+^ and 0.02 mM PQQ and left in incubation for 15 min at 4°C. BUD-ELMs, washed once with 0.01 M PBS, were incubated with reconstituted and non-reconstituted cell lysates for 3.5 h at 4°C and then washed three times with 0.01 M PBS. A small piece of functionalized BUD-ELMs (Fig. S11) was used for the colorimetric assay to detect GDH activity.

### Colorimetric test to detect Glucose Dehydrogenase (GDH) activity

The activity of GDH functionalized material was quantified with modified colorimetric 2,6-dichlorophenol (DCPIP) assay^35^. A reagent solution of 48 mL MOPS buffer (10 mM, pH 7.0, 47 mL), 1mL DCPIP (20 mg dissolved in 5 mL of DI water), and 1 mL phenazine methosulfate (PMS) (45 mg dissolved in 5 mL of DI water) were prepared. Analytical samples were mixed with the reagent to 190 µL. The reaction was initialized by adding 10 µL glucose (1M). Glucose consumption was correlated to the consumption of DCPIP (2:1 ratio), which was quantified colorimetrically by absorption at 600 nm. A representative curve is shown in Fig. S12. The molar absorption coefficient of DCPIP at 600 nm was determined at 21 mM-1cm-1. All the activity assays were performed in a 10 mM MOPS buffer (pH 7.0), including 50 mM glucose, 558 µM PMS, and 262 µM DCPIP at room temperature.

---

**Fig. S1.**
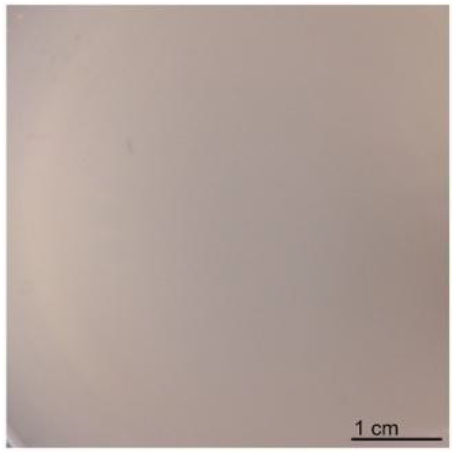
The wild type strain does not form macroscopic aggregates under our growth conditions. Representative image of wild type strain (Mfm126) grown under standard conditions.

**Fig. S2.**
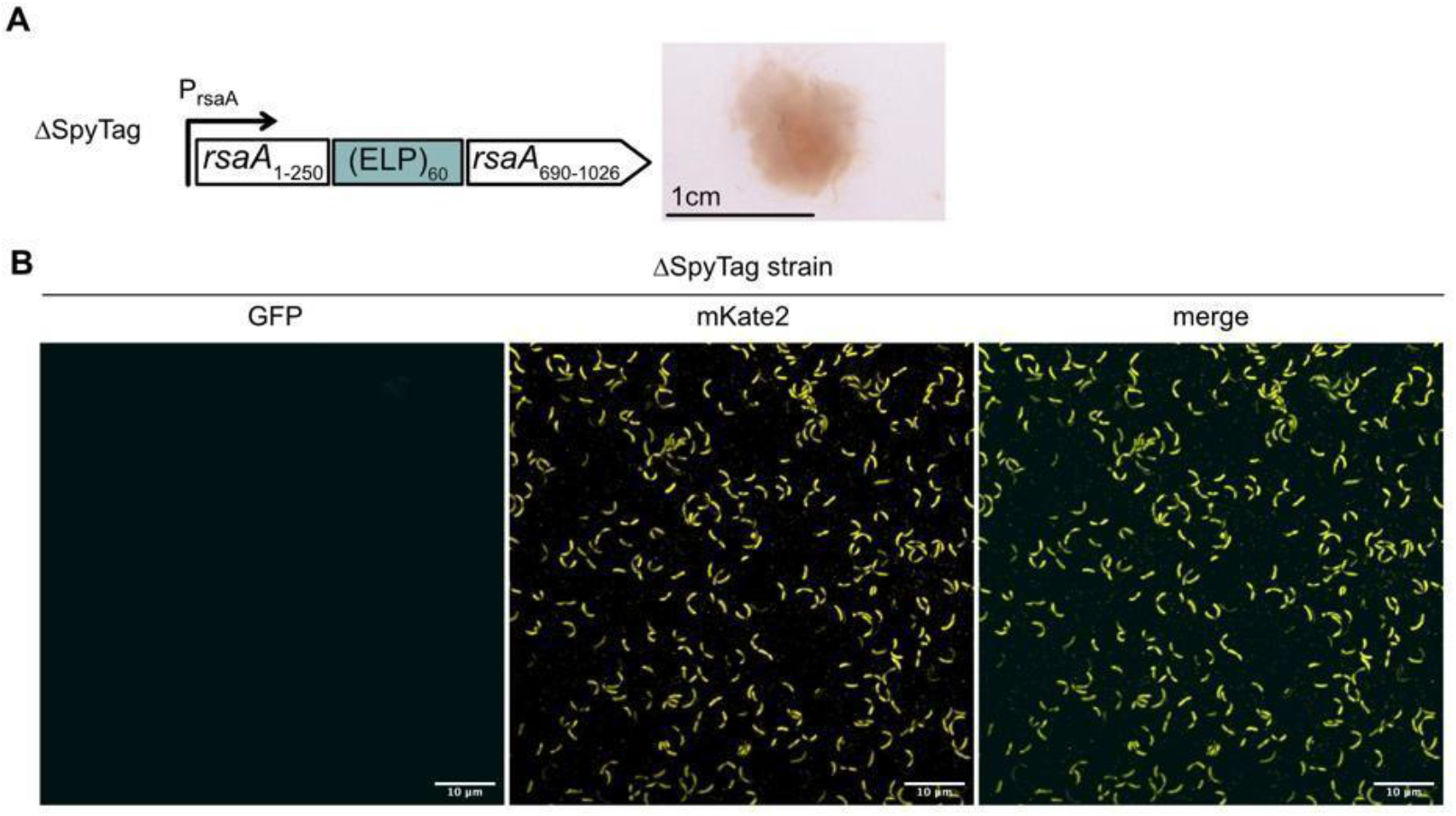
The *ΔSpyTag* strain does not bind SpyCatcher-GFP. (A) Genetic constructs replacing the native *rsaA* gene in the *ΔSpyTag* strain (left panel). Image of corresponding BUD-ELM (right panel) which is indistinguishable from the BUD-ELM containing the SpyTag. (B) Representative image of the *ΔSpyTag* single cells (yellow channel, mKate2) incubated with SpyCatcher-GFP (cyan channel, GFP), showing no significant SpyCatcher-GFP-cell binding.

**Fig. S3.**
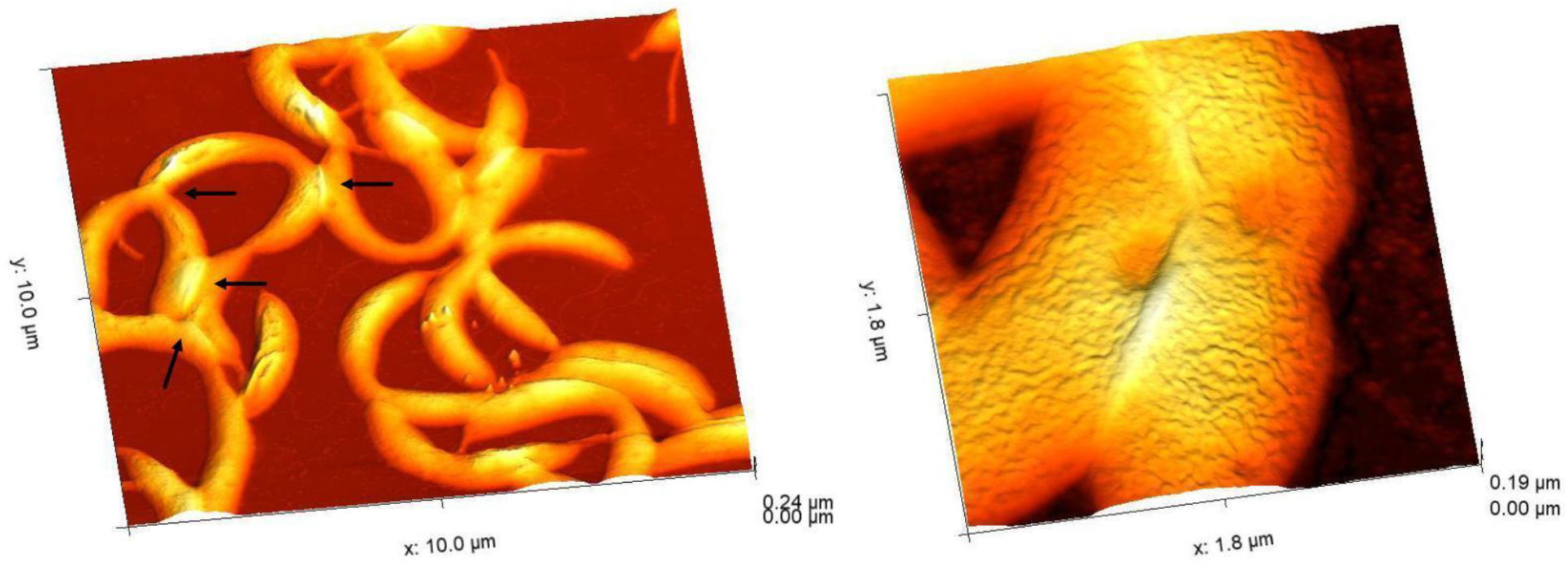
AFM images show cell-cell interactions in the BUD-ELM strain. Representative AFM images at different magnifications of BUD-ELM strain single cells revealing the presence of cell-cell interactions. The junctions of interacting cells show soft material accumulation. BUD-ELM cells also appear with the characteristic brush-like structure of the cell surface.

**Fig. S4.**
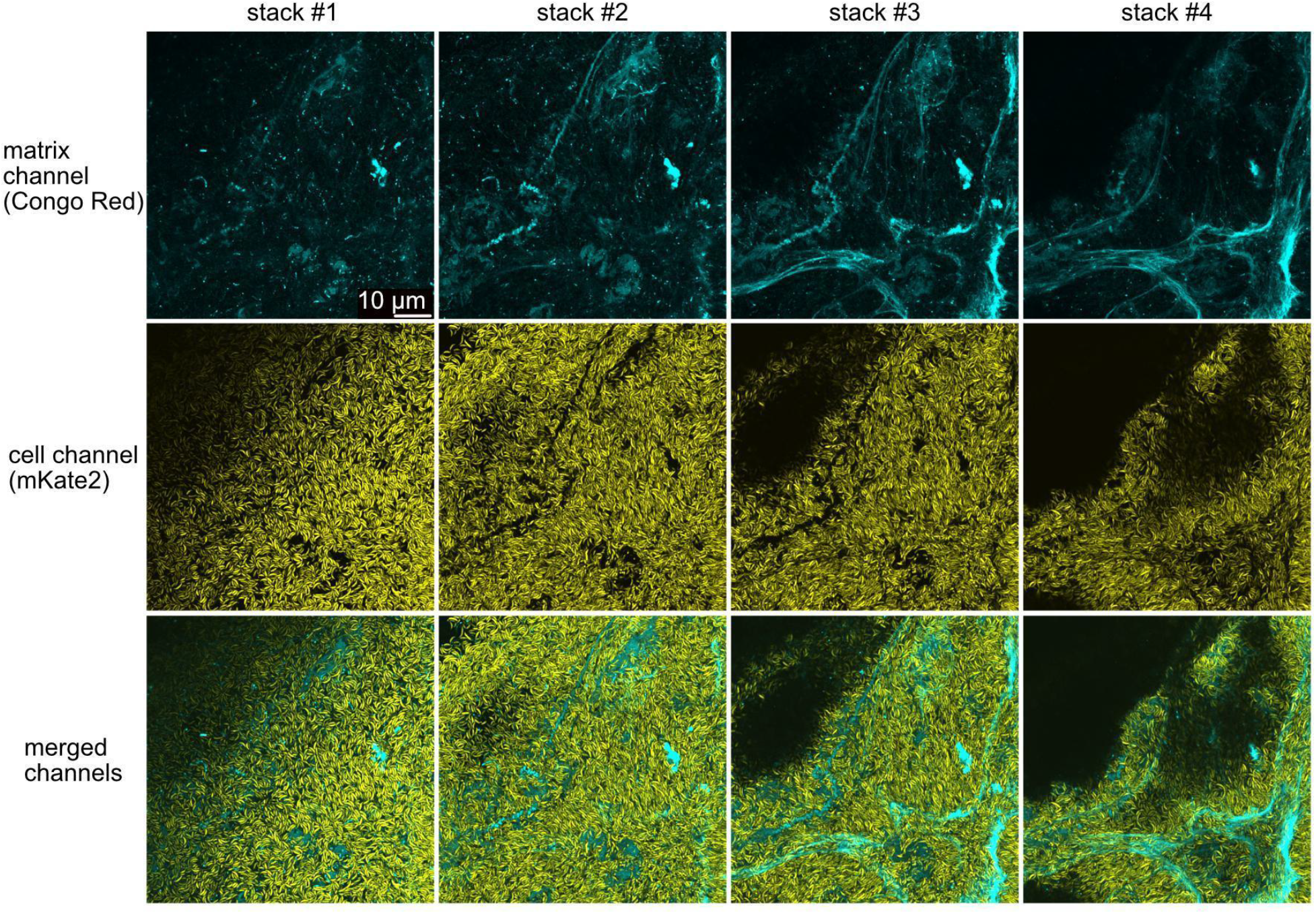
The matrix is surrounded by cells within BUD-ELMs. Representative confocal microscopy stack images of the same BUD-ELM section. They show that the protein matrix, stained with Congo Red, is fully enclosed by cells. Scale bar applies to every image.

**Fig. S5.**
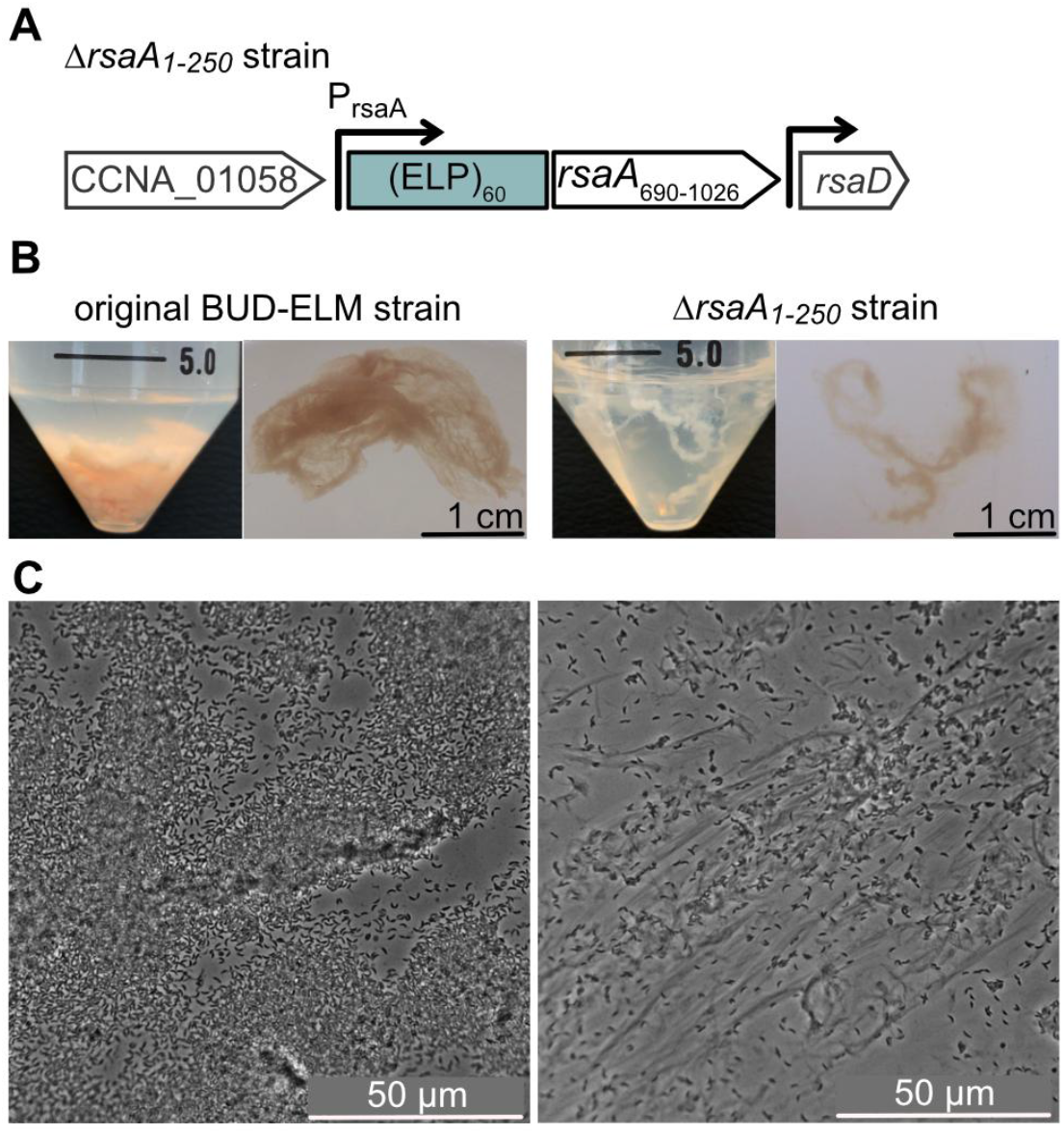
*ΔrsaA_1-250_* strain does not form a cell-rich BUD-ELM. **(A)** Synthetic construct replacing the native *rsaA* gene in the *ΔrsaA_1-250_* strain, showing the missing N-terminal cell anchor domain of the BUD protein. **(B)** Representative images of BUD-ELMs (left panel) and material produced by the *ΔrsaA_1-250_* strain (right panel) growing under standard conditions. The latter material has a distinct morphology (less cohesive) and color (lighter, white) compared to the original BUD-ELM. The material produced by the *ΔrsaA_1-250_* strain appears also more buoyant. (**C**) Optical microscopy of the two materials show a clear difference in cell content, much higher in the original BUD-ELM. This is in agreement with the fact that *ΔrsaA_1-250_* cells do not display the aggregating protein, and therefore cannot specifically adhere to the extracellular protein matrix.

**Fig. S6.**
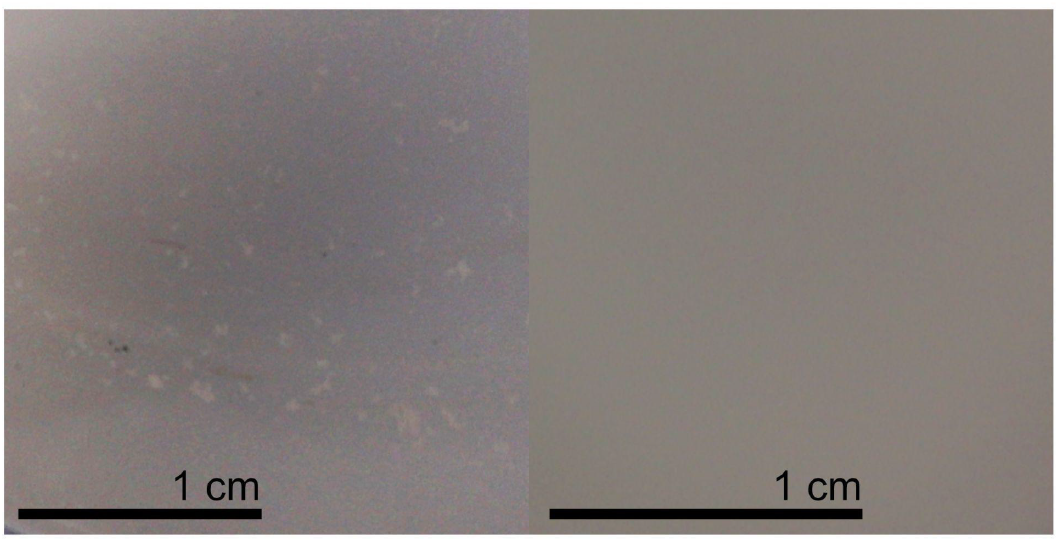
Disruption of the liquid-air interface, through antifoam addition, prevents the formation of large BUD-ELMs. BUD-ELM strain grown under standard conditions with the addition of 0.04% antifoam showed small pieces of material at the bottom of the flask (left). No visible pellicle was observed at the air-water interface (right).

**Fig. S7.**
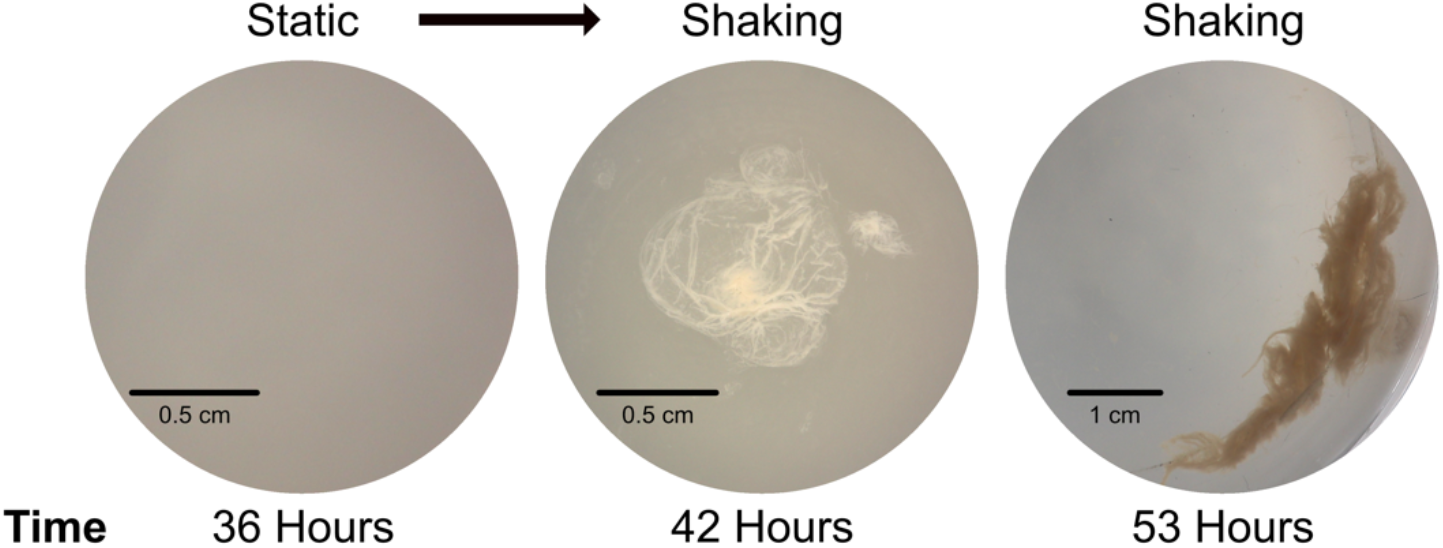
Static-Grown Cultures form BUD-ELM in response to shaking. BUD-ELM strain grown in static conditions showed no material formation after 36 h (left image). When shaking was applied to cultures, pellicle (middle image) and final material (right image) formation were observed within 6 and 11 h, respectively.

**Fig. S8.**
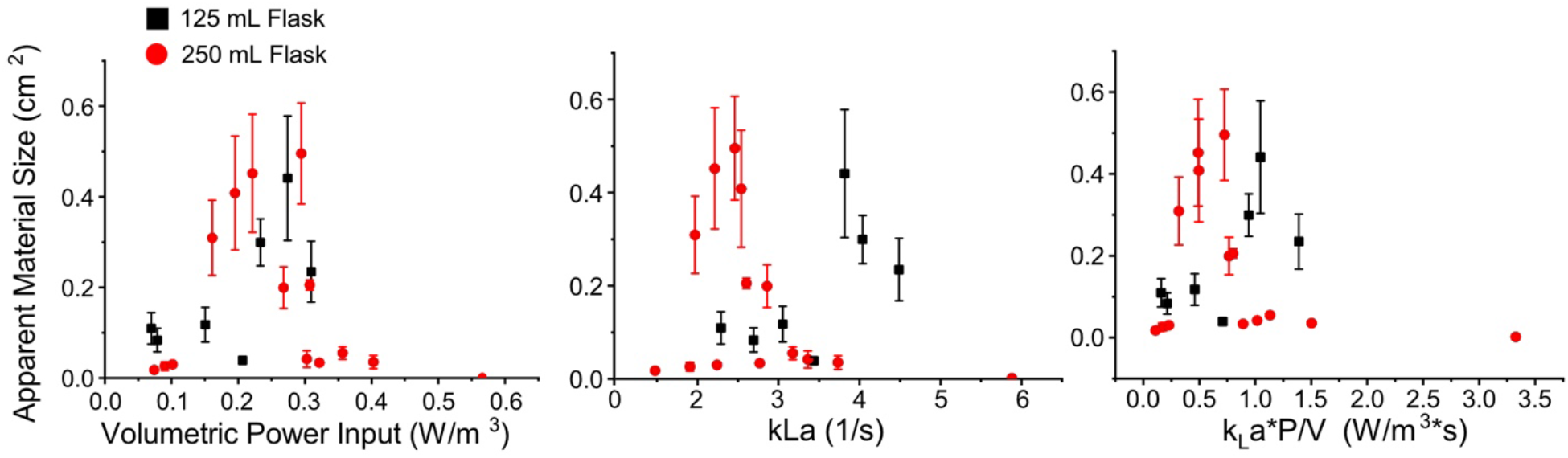
Relationship between the major components of the Modified Volumetric Power and Material Size. Flat surface area of BUD-ELMs was plotted against the Volumetric Power input (left), k_L_a (middle), and the product of the two (right). None of these relationships provide a consistent metric for predicting large BUD-ELM size across flask sizes. Error bars represent standard error of at least three samples. X-axis label and legend apply to every graph.

**Fig. S9.**
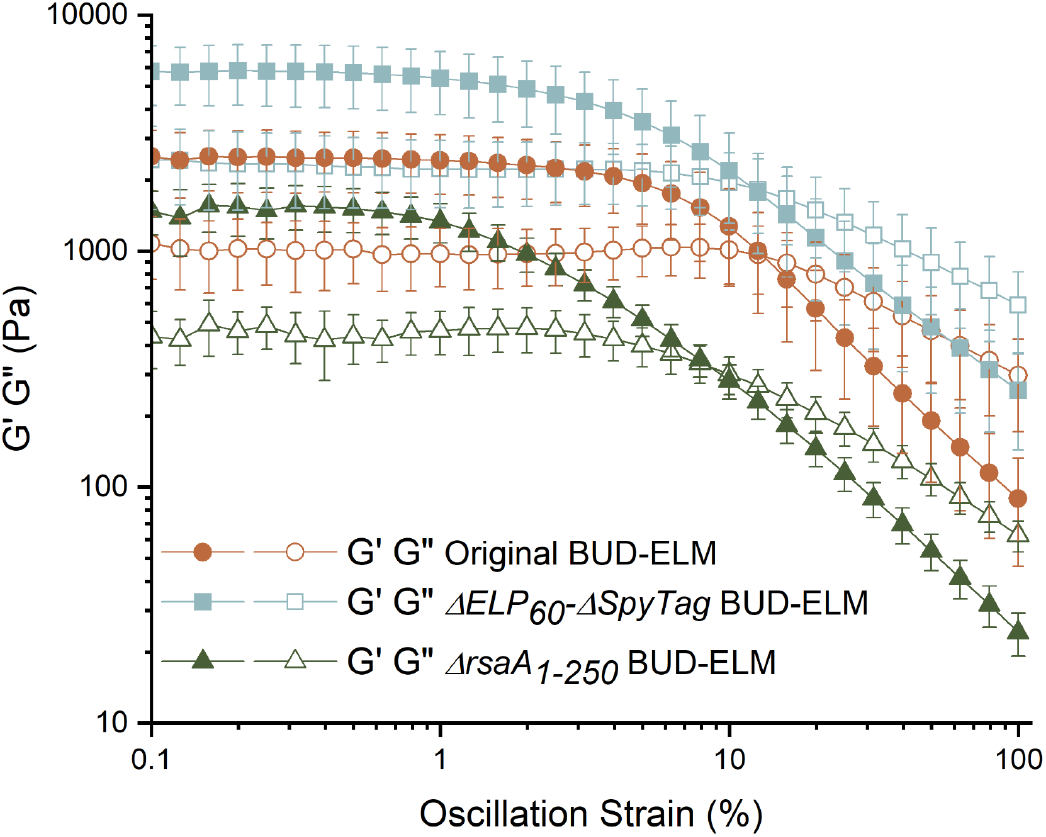
Stress-strain curves of BUD-ELMs. Strain sweep measurements were acquired from 0.1% to 100% strain amplitude at a constant frequency of 3.14 rad/s. Error bars represent 95% confidence intervals of at least five samples. From the amplitude sweep curves, we identified the linear viscoelastic region of the three BUD-ELMs and set the strain used to collect frequency sweep data (Fig. S10) to 0.35%.

**Fig. S10.**
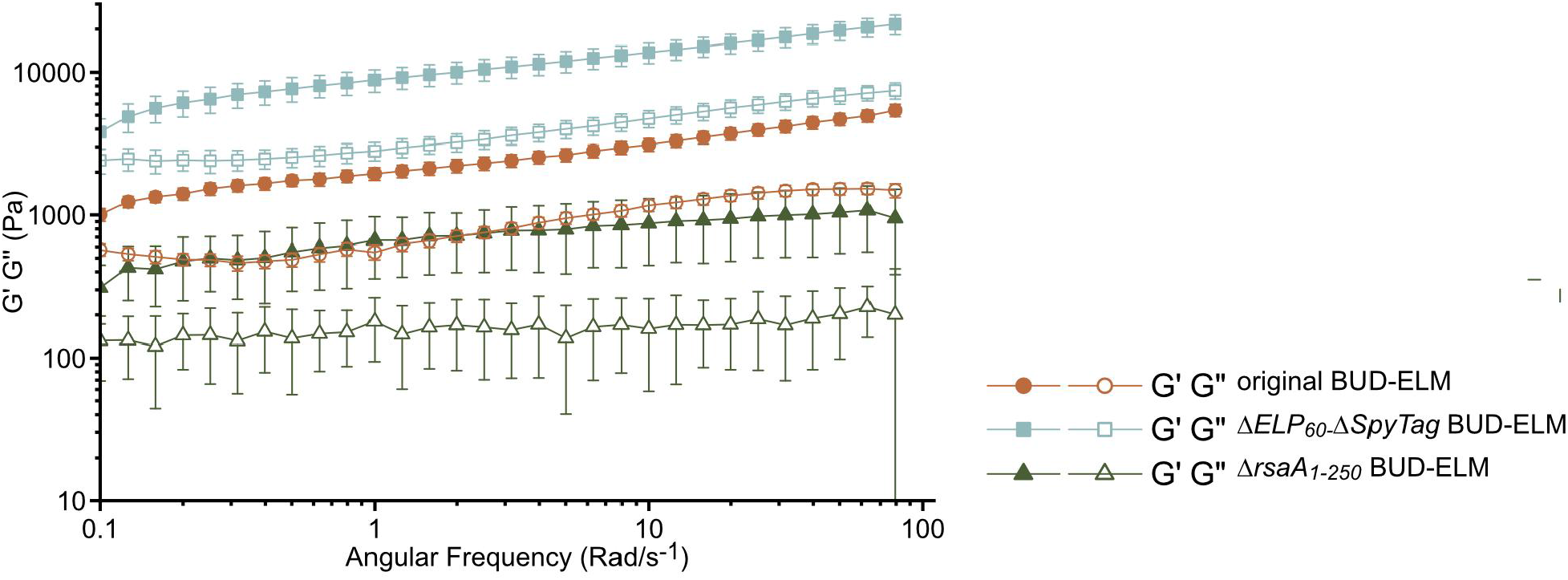
Frequency sweep curve. Frequency sweep measurements were acquired from 0.1 rad/s to 100 rad/s at a constant strain amplitude of 0.35%. Error bars represent 95% confidence intervals of at least five samples.

**Fig. S11.**
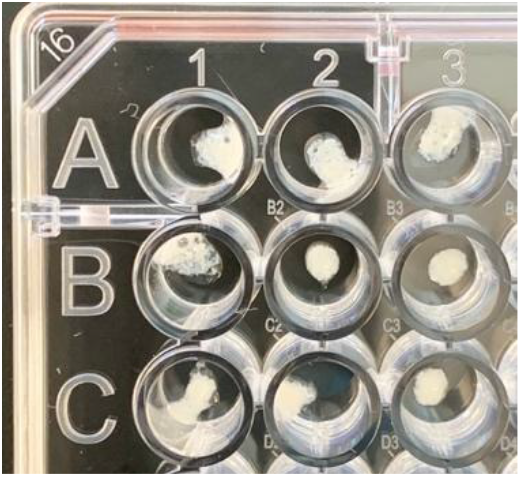
Representative amount of BUD-ELM used the GDH activity colorimetric test. Each well contains a comparable amount of functionalized BUD-ELM.

**Fig. S12.**
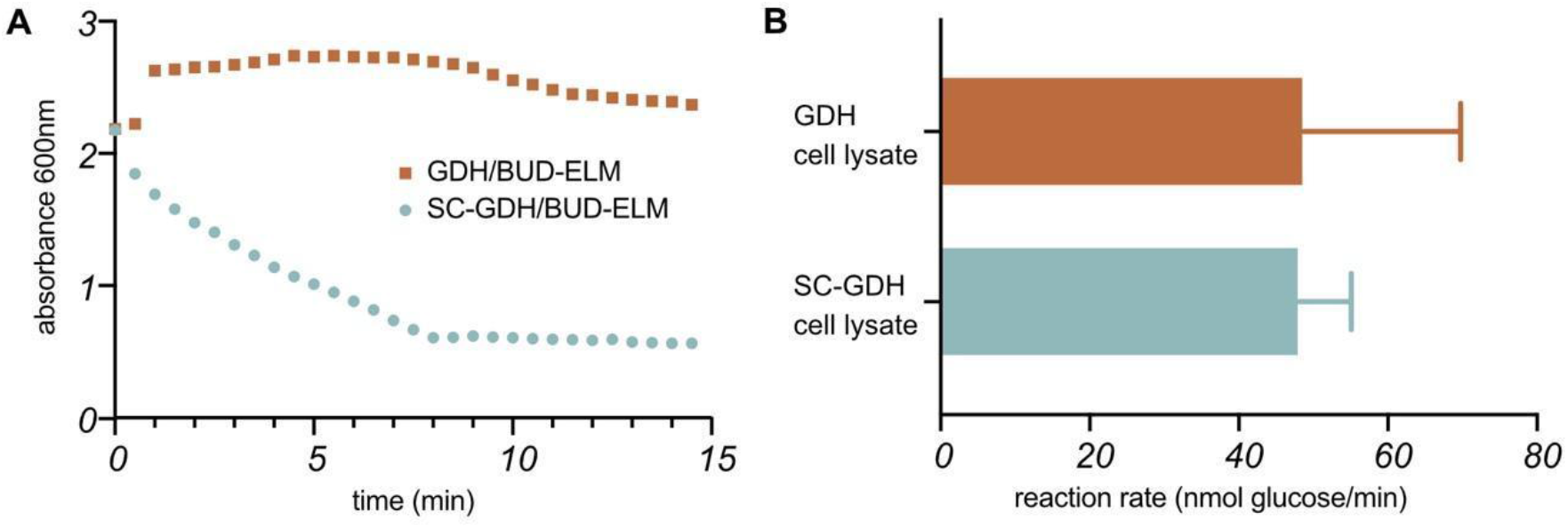
Reaction rate of the cell lysate extracted from strains producing SpyCatcher-holo-GDH and holo-GDH. (A) Representative kinetic curve of the enzymatic conversion of 2,6-dichlorophenol–indophenol (DCPIP) by BUD-ELM incubated with SpyCatcher-holo-GDH (circles) or holo-GDH (squares). (B) Both strain lysate containing SpyCatcher-holo-GDH and holo-GDH show enzymatic activity. Error bars represent standard error of three samples.

**Table S1.** List of Strains used in this work.

| Strain Name | Strain Designation | Source | genotype |
| --- | --- | --- | --- |
| wild type | MFm126 | A. Persat *et al*<sup>35</sup> | NA1000 $\Delta sapA :: Pxyl mKate2$ |
| $\Delta rsaA$ | MFm133 | M. Charrier *et al*<sup>20</sup> | NA1000 $\Delta rsaA :: Spec$ |
| $\Delta rsaA_{1-250}$ | MFm152 | M. T. Orozco-Hidalgo *et al*<sup>21</sup> | NA1000 $\Delta sapA :: Pxyl-mKate2$,<br>$\Delta rsaA(1-689) :: ELP_{60}$ |
| original<br>BUD-ELM | RCC002 | This work | NA1000 $\Delta sapA :: Pxyl mKate2$,<br>$\Delta rsaA(251-689) :: ELP_{60}-SpyTag$,<br>(Fig. 1B). |
| $\Delta ELP_{60}\Delta SpyTag$ | RCC004 | This work | NA1000 $\Delta sapA :: Pxyl mKate2$,<br>$\Delta rsaA(251-689)$, (Fig. 1G). |
| $\Delta SpyTag$ | RCC005 | This work | NA1000 $\Delta sapA :: Pxyl mKate2$,<br>$\Delta rsaA(251-689) :: ELP_{60}$, (Fig. 1F). |

**Table S2.** List of Plasmids used in this work.

| Plasmid | Source | Description |
| --- | --- | --- |
| pSMCAF008 | This work | Integration plasmid for strain RCC002. *rsaA-N-terminus* (800bp), *(ELP)<sub>60</sub>*, *SpyTag*, *rsaA-C-terminus* (800bp) |
| pSMCAF017 | This work | Integration plasmid for strain RCC004. *rsaA-N-terminus* (800bp), *rsaA-C-terminus* (800bp) |
| pSMCAF018 | This work | Integration plasmid for strain RCC005. *rsaA-N-terminus* (800bp), *(ELP)<sub>60</sub>*, *rsaA-C-terminus* (800bp) |
| pSMCAF015 | This work | *pAra*, *HisTag-Tev-GFP*; L-arabinose-induced expression of GFP, used for protein purification (Fig. 1D). |
| pSMCAF016 | This work | *pAra*, *HisTag-Tev-SpyCatcher-GFP*; L-arabinose-induced expression of SpyCatcher-GFP, used for protein purification (Fig. 1D and Fig. 2A). |
| pSMCAF029 | This work | *pAra*, *HisTag-Tev-SpyCatcher-GDH*; L-arabinose-induced expression of SpyCatcher-GDH, used for BUD-ELM functionalization (Fig. 4F). |
| pSMCAF032 | This work | *pAra*, *HisTag-Tev-GDH*; L-arabinose-induced expression of GDH, used for BUD-ELM functionalization (Fig. 4F). |
| KR-12 | Provided by Kathleen R. Ryan | *pTac*, *GFP-mut3*. IPTG-induced expression of GFP-mut3 in *C. crescentus*. (Fig. 2B – Congo Red staining). |

**Table S3.** List of Growth conditions (flask volume, shaking speed and culture volume) used to generate the model in Fig. 3D.

| Flask Volume (mL) | Shaking Speed (rpm) | Culture Volume (mL) |
| --- | --- | --- |
| 125 | 150 | 25 |
| 125 | 150 | 30 |
| 125 | 200 | 30 |
| 125 | 225 | 25 |
| 125 | 225 | 30 |
| 125 | 250 | 25 |
| 125 | 250 | 30 |
| 250 | 150 | 50 |
| 250 | 150 | 60 |
| 250 | 150 | 80 |
| 250 | 200 | 60 |
| 250 | 200 | 80 |
| 250 | 200 | 80 |
| 250 | 225 | 50 |
| 250 | 225 | 60 |
| 250 | 225 | 80 |
| 250 | 250 | 30 |
| 250 | 250 | 50 |
| 250 | 250 | 60 |
| 250 | 250 | 70 |
| 250 | 250 | 75 |
| 250 | 250 | 80 |

